# MADS31 supports female germline development by repressing the post-fertilization program in cereal ovules

**DOI:** 10.1101/2022.12.05.519106

**Authors:** Xiujuan Yang, Gang Li, Jin Shi, Laura G. Wilkinson, Matthew K. Aubert, Kelly Houston, Neil J. Shirley, Lucia Colombo, Matthew R. Tucker

**Affiliations:** Waite Research Institute, School of Agriculture, Food and Wine, The University of Adelaide, Urrbrae, South Australia, 5064, Australia; Nanjing Agricultural University, Nanjing, 210095, China; Shanghai Jiao Tong University, Shanghai, 200240, China; The James Hutton Institute, Invergowrie, Dundee, DD2 5DA, Scotland, UK; Department of Biosciences, Università degli Studi di Milano, Via Celoria 26, 20133 Milan, Italy; Australian Grain Technologies, Northam, Western Australia, 6401, Australia

## Abstract

The female germline of flowering plants develops within a niche of somatic ovule cells, also referred to as the nucellus. How niche cells maintain their own somatic developmental program, yet support the development of adjoining germline cells, remains largely unknown. Here we report that MADS31, a conserved MADS-box transcription factor from the B-sister subclass, is a potent regulator of niche cell identity in barley. MADS31 is preferentially expressed in nucellar cells directly adjoining the germline, and loss-of-function *mads31* mutants exhibit deformed and disorganized nucellar cells, leading to impaired germline development and partial female sterility. Molecular assays indicate that MADS31 encodes a potent transcriptional repressor, repressing genes in the ovule that are normally active in the seed. One prominent target of MADS31 is *NRPD4b*, a seed-expressed component of RNA polymerase IV/V that is involved in gene silencing via RNA directed DNA methylation. *NRPD4b* is directly repressed by MADS31 *in vivo* and is de-repressed in *mads31* ovules, while overexpression of *NRPD4b* recapitulates the *mads31* ovule phenotype. This coincides with specific changes in histone methylation and is consistent with *NRPD4b* being directly repressed by MADS31 to maintain ovule niche functionality. Our findings reveal a new mechanism by which somatic ovule tissues maintain their own identity before transitioning to the post-fertilization program.

## Introduction

In seed-bearing plants, ovules are a complex mixture of somatic (diploid) and gametophytic (haploid) tissues that perform distinct roles during reproduction. Prominent somatic tissues include a distal nucellus, which gives rise to the female germline and gametophyte; the central chalaza, which gives rise to the protective integuments and seed coat; and the proximal funiculus, which connects the ovule to a supply of maternal nutrients (Rudall, 2021). The formation and differentiation of these tissues creates a scaffold for seed development. During early development, the ovule initially enters a generative phase, during which the germline is established within a pool of somatic nucellus cells. Specifically, a single nucellus cell initiates megasporogensis and differentiates into a megaspore mother cell (MMC). This cell divides by meiosis to give rise to four haploid megaspores, one of which typically undergoes three rounds of mitosis to form a single female gametophyte (embryo sac), which contains an egg cell, two synergid cells, a central cell, and multiple antipodal cells (Bowman et al., 1994). As the ovule matures, the integuments form a protective coat around the embryo sac, and the nucellus show signs of collapse in preparation for fertilization and seed initiation.

Also referred to as the megasporangium, the nucellus represents a niche that supports germline development from initiation until fertilization (Rudall, 2021). This function appears to be independent of the number of cells within the nucellus, which varies from relatively few cells in dicotyledonous Arabidopsis, to many in monocotyledonous cereals such as rice and barley (Endress, 2011). Genetic studies indicate this niche integrates multiple regulatory pathways to support germline initiation (Pinto et al., 2019). Defects at this generative stage range in severity from subtle to extreme. For example, *sporocyteless* mutants produce a nucellus-like tissue but fail to initiate a germline (Yang et al., 1999); mutations in the RNA-directed DNA Methylation (RdDM) machinery such as RDR6 and AGO9 result in extra nucellar cells adopting features of germline-like cells (Olmedo-Monfil et al., 2010); and weak PIN1 (Ceccato et al., 2013) or null RGTB1 (Rab Geranylgeranyl Transferase Beta Subunit) mutants where PIN1 function is compromised produce a functional megaspore that fails to initiate gametogenesis (Rojek et al., 2021).

The temporal and spatial coordination of these pathways has remained unclear, although recent progress suggests that the D-class MADS-box gene SEEDSTICK (STK) acts upstream of RdDM pathways in the nucellus to spatially regulate SPL expression, which in turn promotes auxin signalling (Mendes et al., 2020). During later stages of gametophyte development, auxin signalling also regulates nucellus degeneration, which occurs concomitantly with rapid expansion of the embryo sac (Wang et al., 2021). In cereal species, nucellar degeneration continues after fertilization and involves a range of programmed cell death-related genes including the novel *Jekyll* protein (Radchuk et al., 2006). This coincides with the establishment of transfer tissues that provide maternally-derived nutrition to the endosperm (Daneva et al., 2016; Wang et al., 2020). In most seed plants, transfer tissues differentiate from the nucellus or integuments (Xu et al., 2016; Yin and Xue, 2012), and in wheat and barley, the last vestige of the nucellus is referred to as the nucellar projection. Taken together, maternal regulation via nucellar tissue is essential for multiple stages of sexual reproduction, however the transcriptional drivers of nucellus maintenance, degeneration and differentiation remain largely unknown.

Along with STK, the plant-specific type II MIKC C-, D-, and E-class MADS box genes control ovule initiation (Favaro et al., 2003) as well tissue differentiation during ovule and seed development (Pinyopich et al., 2003; Brambilla et al., 2007). The subclass B sister (Bsis) MADS-box proteins, also from the type II MIKC family, are mainly expressed in female reproductive organs (Becker et al., 2002; Frohlich, 2003). The Arabidopsis genome contains two Bsis genes, *TRANSPARENT TESTA16 (TT16)/ARABIDOPSIS BSISTER MADS* (*ABS*) and *GORDITA* (*GOA*)/*AGL63. ABS* promote nucellus degradation (Xu et al., 2016), specifies the endothelium via interaction with *SEEDSTICK* (Mizzotti et al., 2012), and supports deposition of pigments in the maturing integuments and seed coat (Nesi et al., 2002). *GOA*, a paralog of *ABS*, is more widely expressed in ovules, seeds and fruits, contributing to fruit growth and integument/seed coat development (Prasad et al., 2010; Erdmann et al., 2010). However, neither *abs* nor *goa* show significant ovule abortion, indicating their dispensable roles in determining ovule fertility. In orchids, Bsis *PeMADS28* is expressed in the nucellus and integuments, and can rescue the Arabidopsis *abs* mutant (Shen et al., 2021). In cereals, among three subclades of Bsis (MADS29, MADS30 and MADS31; Yang et al., 2012), MADS29 is vital to seed formation. The *MADS29* gene is expressed in the nucellus the residual nucellar projection during seed development, and loss-of-function mutations cause seed abortion, a consequence of no endosperm development (Yin and Xue, 2012; Yang et al., 2012; Lee et al., 2013; Shoesmith et al., 2021). Rice MADS30 regulates plant architecture and is not involved in female reproduction, a likely product of evolutionary neofunctionalization of this member of the MADS-box family (Schilling et al., 2015). A role for the third member of the Bsis family in cereals, MADS31, has yet to be elucidated.

Here we show that *MADS31* is preferentially expressed in the barley nucellus and functions to maintain nucellus development. Furthermore, we provide evidence to suggest that MADS31 represses transcription of key genes involved in post-fertilization development, including RdDM and cell death pathways, therefore maintaining the integrity of the ovule nucellus. Our findings provide new insight regarding the maintenance of somatic ovule cells, and their role in coordinating female germline development.

## Results

### *MADS31* expression is preferentially expressed in the inner nucellus of barley ovules

Previously we established a tissue-specific transcriptome atlas for the barley ovule (Wilkinson, 2019a; Yang et al., 2022). Cells were captured from the nucellus, integuments, ovary wall, and embryo sac at different stages of ovule development before RNA was extracted and sequenced (Fig. 1a). Genes that were preferentially expressed in the nucellus were selected (examples are shown in Extended Data Fig. 1). One of these, a B-sister (Bsis) class MADS-box gene named *MADS31*, was abundant in the nucellus and showed lower expression in the integuments and embryo sac (Fig. 1b). Notably, *MADS31* was the most abundant Bsis in the ovule relative to its paralogs *MADS29* and *MADS30* (Extended Data Fig. 1). Real-time PCR showed *MADS31* was specifically expressed in the pistil, and was absent in vegetative tissues or other floral organs (Extended Data Fig. 2a). In the pistil, *MADS31* transcripts increased as the ovule developed from a primordial (Ov1) (Wilkinson et al., 2019b) to mature female gametophyte stage (Ov9a) then decreased at anthesis (Ov9a and Ov10) (Extended Data Fig. 2b). We examined tissue specificity of *MADS31* transcript and protein using mRNA *in situ* hybridization, and a transgenic line expressing a translational fusion of MADS31 to enhanced green fluorescent protein (eGFP) under the control of its native promoter (*pMADS31*). At Ov2 stage, when the primary germline cell distinguishes itself from adjoining nucellar cells, *MADS31* transcript and protein begin to accumulate in the nucellar cells adjacent to the archesporial cell. As the archesporial cell develops into the megaspore mother cell (MMC; Ov3) and undergoes meiosis (Ov4) and mitosis (Ov7/8) to form the embryo sac, MADS31 expression gradually spreads to two or three cell layers of the nucellus, which surround the germline, in a zone hereafter referred to as the inner nucellus. In ovules approaching maturity (Ov9b/10), MADS31 was also observed in the outer nucellus, which surrounds the inner nucellus (Fig. 1c). Besides localization to the nucellus, MADS31 was also weakly detected within part of the inner integument adjacent to the nucellus apex and ovary wall (Fig. 1c and Extended Data Fig. 2c). Intriguingly, *MADS31* transcripts were detected in the embryo sac by LCM-RNA-sequencing as well as *in situ* hybridization, however no GFP signal could be observed within the germline at any stage. This suggests MADS31 protein is restricted to somatic maternal cells, and its expression distinguishes the inner from the outer nucellus until the maturity of ovule.

**Fig. 1:**
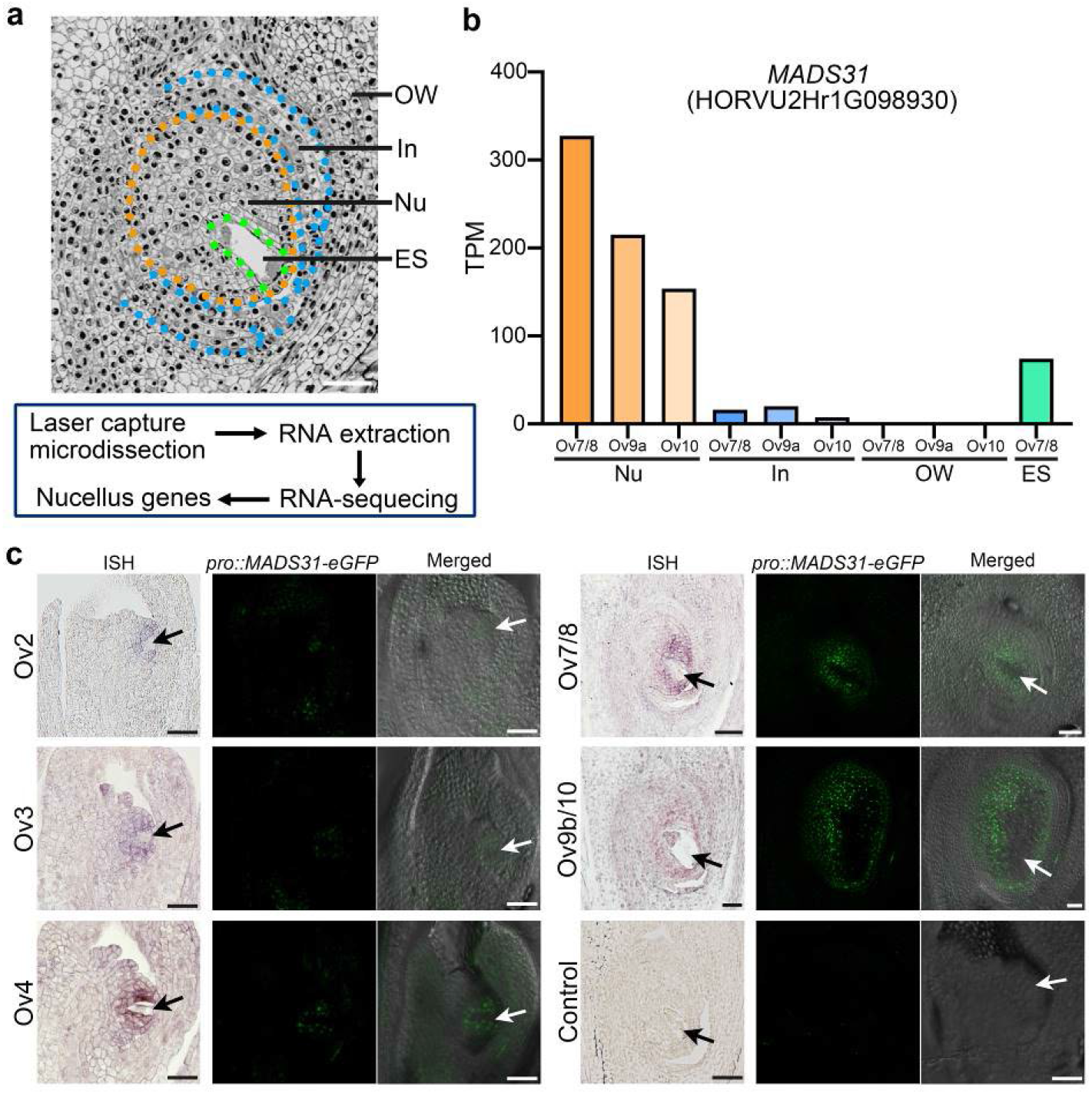
*MADS31* is expressed in a restricted niche in the nucellus of barley ovule. **a**, The procedure of tissue-specific transcriptome profiling. OW, ovary wall; In, integuments; Nu, nucellus; ES, embryo sac/female gametophyte. Scale bar, 50 μm. **b**, *MADS31* expression in ovule tissues by RNA-sequencing. TPM, transcripts per million. Ov7/8, female gametophyte (FG) mitosis stage; Ov9a, mature FG stage; Ov10, FG at anthesis stage. **c**, Accumulation of *MADS31* transcripts and MADS31 proteins shown by in situ hybridization and vibratome sections of *pro::MADS31-eGFP* plants, respectively. Black and white arrows indicate the megaspore mother cell (MMC) or female gametophyte. Ov2, archesporial cell stage; Ov3, MMC stage; Ov4, meiosis stage. Ov9b, FG expansion stage. Sense probe and non-transgenic plants are served as negative controls. Scale bars, 50 μm.

### Inner nucellus and embryo sac development are impaired in *mads31* ovules

To investigate the role of *MADS31* in nucellus development, we used CRISPR/Cas9 gene editing (Ma et al., 2015) to generate loss-of-function alleles in barley cv. Golden Promise. Multiple alleles were identified, including a range of insertions and deletions that compromise production of MADS31 protein (Extended Data Fig. 3a). Putative null alleles showed a similar reduction in seed set (Extended Data Fig. 3b), and one of these was selected *(mads31#12)* for detailed investigation, hereafter referred to as *mads31*. Compared with wild-type spikes, *mads31* spikes produced approximate 70% less seeds, verified over two successive generations (Extended Data Fig. 3c). Clearing of mature pistils showed that *mads31* ovules produce smaller embryo sacs, approximately half size of that in wild-type (Extended Data Fig. 3d, e). At anthesis, *mads31* pistils appeared normal in terms of ovary size, style and stigma formation. Furthermore, mature anthers from *mads31* spikelets were yellow and produced viable pollen, indistinguishable from wild-type (Extended Data Fig. 3e). These results suggest that reduced seed set in *mads31* is likely to be a consequence of defective ovule development.

We further examined wild-type and *mads31* ovules using histological sectioning. In wild-type ovules at Ov2 stage, one cell beneath the nucellus apex was enlarged and trapezoid in shape, indicative of the germline archesporial cell (Fig. 2a). The hypodermal nucellar cells that normally express *MADS31* (Fig. 1c) adjacent to the germline were rectangular and aligned uniformly (Fig. 2a). Immunolabelling showed that these cells are labelled by the LM19 antibody, which recognises de-esterified pectin in the cell wall, and is a hallmark of cell stiffness (Fig 2a; Yang et al., 2022). Based on their position close to the germline, we defined these cells as the inner nucellus. At the same stage in *mads31* ovules, the inner nucellar cells were deformed, rounded and disorganized relative to wild-type in over half of the observed ovules. Additionally, the accumulation of de-esterified pectin was heavily reduced (Fig. 2a).

**Fig. 2:**
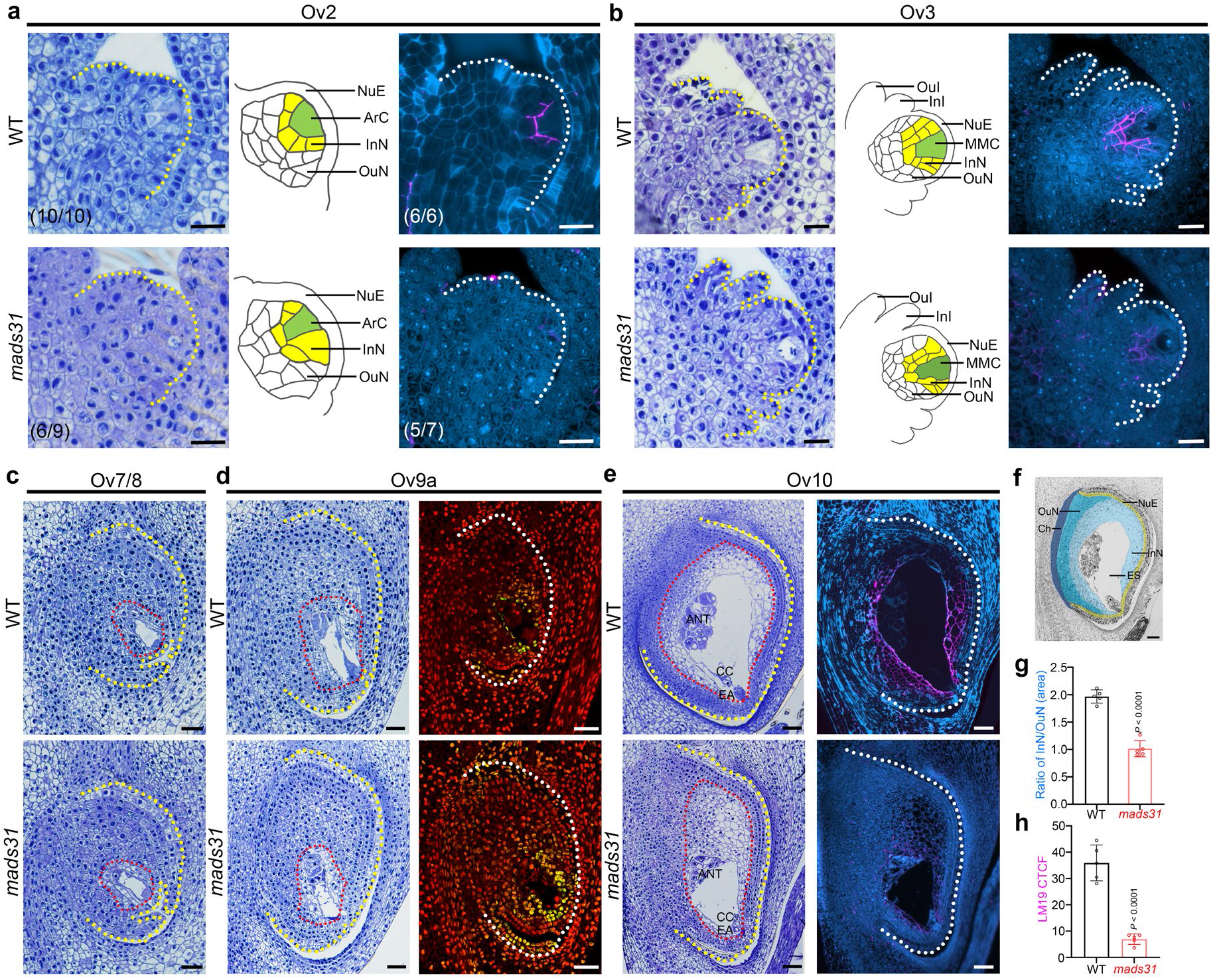
Inner nucellus and embryo sac development are impaired in *mads31* ovules. **a, b,** Early stages (Ov2 and Ov3) of ovule development in WT and *mads31*. Left, toluidine blue stained longitudinal sections of WT and *mads31* ovules. Middle, diagrams of cell arrangement in the nucellus. Green, the germ cell; yellow, inner nucellus. Right, LM19 labeled demethylesterified pectin in the cell walls of inner nucellus. OuI, outer integumens; InI, inner integuments; NuE, nucellus epidermis; ArC, archesporial cell, MMC, megaspore mother cell; InN, inner nucellus, OuN, outer nucellus. Scale bars, 25 μm. **c, d,** Middle stages (Ov7/8 and Ov9a) of ovules in WT and *mads31*. In **d,** left, toluidine blue stained longitudinal sections; right, TUNEL assay in WT and *mads31* ovules. Scale bars, 50 μm. **e,** The final stage (Ov10) of ovules in WT and *mads31*. Left, toluidine blue stained longitudinal sections; right, LM19 labeled demethylesterified pectin in the cell walls of inner nucellus. Yellow and white dashed lines indicate the ovule within the carpel and red dashed lines indicate the region of inner nucellus. ANT, antipodal cells; CC, central cell; EA, egg apparatus. **f, g,** colored regions of nucellus from semi-thin section and statistical analysis for the ratio of inner nucellus area versus outer nucellus. ES, embryo sac/female gametophyte; Ch, chalaza. Scale bar, 50 μm. The data are shown as mean±s.d.; *n = 5* ovules. **h,** Corrected total cell fluorescence (CTCF) of LM19 immunosignals in the nucellus of wild-type and *mads31* ovules. The data are shown as mean ±s.d.; *n* = 5 ovules.

At stage Ov3, when the germline differentiates into an MMC in wild-type ovules, the rectangular inner nucellar cells still contained de-esterified pectin in their walls but had divided to form a multi-layered tissue (Fig. 2b). The same region in *mads31* ovules showed different degrees of abnormality and was defined by cells of irregular shape that contained low levels of de-esterified pectin in their cell walls (Fig. 2b and Extended Data Fig. 4a, b). After megaspore selection and the initiation of embryo sac mitosis, the innermost cells of the nucellus in wild-type ovules exhibited mild vacuolation reminiscent of cell degeneration. This vacuolation of inner nucellar cells was more obvious in *mads31* ovules and extended further outwards into the nucellar tissue (Fig. 2c, d and Extended Data Fig. 4c, d). A terminal deoxynucleotidyl transferase dUTP nick end labeling (TUNEL) assay confirmed additional cell death events within the nucellus of *mads31* relative to wild-type (Fig. 2d), suggesting that changes in cell shape and vacuolation in *mads31* might correspond to changes in cell identity and/or viability.

In wild-type plants, female germline maturity is attained at stage Ov10 when the ovule contains a fully expanded embryo sac. The mature embryo sac incorporates a cluster of antipodal cells at the apical pole, a central cell that is located between the apical and basal poles, and an egg apparatus that is located at the basal (micropylar) end. At this stage in wild-type ovules, differences between the inner and outer nucellus become more striking. Inner nucellar cells are enlarged, lack obvious cytoplasm, and their walls are rich in de-esterified pectin, whereas those of the outer nucellus are more rounded, contain dense cytoplasm, and lack de-esterified pectin (Fig. 2e and Extended Data Fig. 4e). In *mads31*, the embryo sac is much smaller (approximately half as large; Extended Data Fig. 3e), containing residue of degenerated antipodal cells or morphologically abnormal antipodal cells with cytoplasmic condensation as a sign of cellular degeneration (Fig. 2e and Extended Data Fig. 4f). The central cell nucleus was sometimes observed to be directly adjacent to the embryo sac wall or present at the basal micropylar end, and the egg apparatus was mildly vacuolated. In the ovule proper, *mads31* exhibited a larger proportion of outer nucellus, and inner nucellus expansion was inhibited, supported by the decreased ratio of area between inner nucellus and outer nucellus in *mads31* ovules (Fig. 2f, g). The outer nucellus cells were small and appeared to have overproliferated, taking up much of the room in the ovule. Also, LM19 immunolabeling showed a clear reduction in de-esterified pectin within the inner nucellus (Fig. 2e, h). Importantly, a small proportion of *mads31* ovules exhibited less severe morphological changes (Extended Fig. 4b, f). Given that *mads31* can still produce a reduced number of viable seeds, we assume these ovules can be fertilized.

Taken together, these results demonstrate that nucellus differentiation and patterning in barley starts at germline initiation and continues until anthesis. Loss of MADS31 function impairs the development of the inner nucellus by altering cell morphogenesis and causing premature cellular degeneration. Although MADS31 protein is absent from the germline, the embryo sac of *mads31* ovules exhibit various defects, consistent with a maternal influence of the inner nucellus on germline development.

### Loss of MADS31 reprograms the ovule via transcriptional activation

To further understand the molecular basis for MADS31 function, we compared the transcriptome of wild-type and *mads31* at several stages. Spikes including ovules at stage Ov2 and Ov3/4 were selected to investigate transcriptional changes during sporogenesis, whereas pistils at stage Ov7/8 stage were selected to investigate changes during gametogenesis. In total, 1,263 differentially expressed genes (DEGs) were identified. At Ov2 stage, only 192 DEGs were found, including 98 up-regulated and 94 down-regulated DEGs. At stages Ov3/4 and Ov7/8 the number of DEGS increased significantly and the majority were up-regulated (67.7% and 61.4%, respectively) (Fig. 3a, Extended Data Fig. 5a and Supplementary Dataset 1), suggesting loss of MADS31 may trigger transcriptional activation. Gene ontology (GO) enrichment analysis indicated that DEGs involved in gene silencing, RNA processing, stress response and cell death control were altered throughout all stages. At stages Ov3/4 and Ov7/8, DEGs related to metabolic processes, transmembrane transport and cell wall remodeling were also enriched (Fig. 3b and Supplementary Dataset 4).

**Fig. 3:**
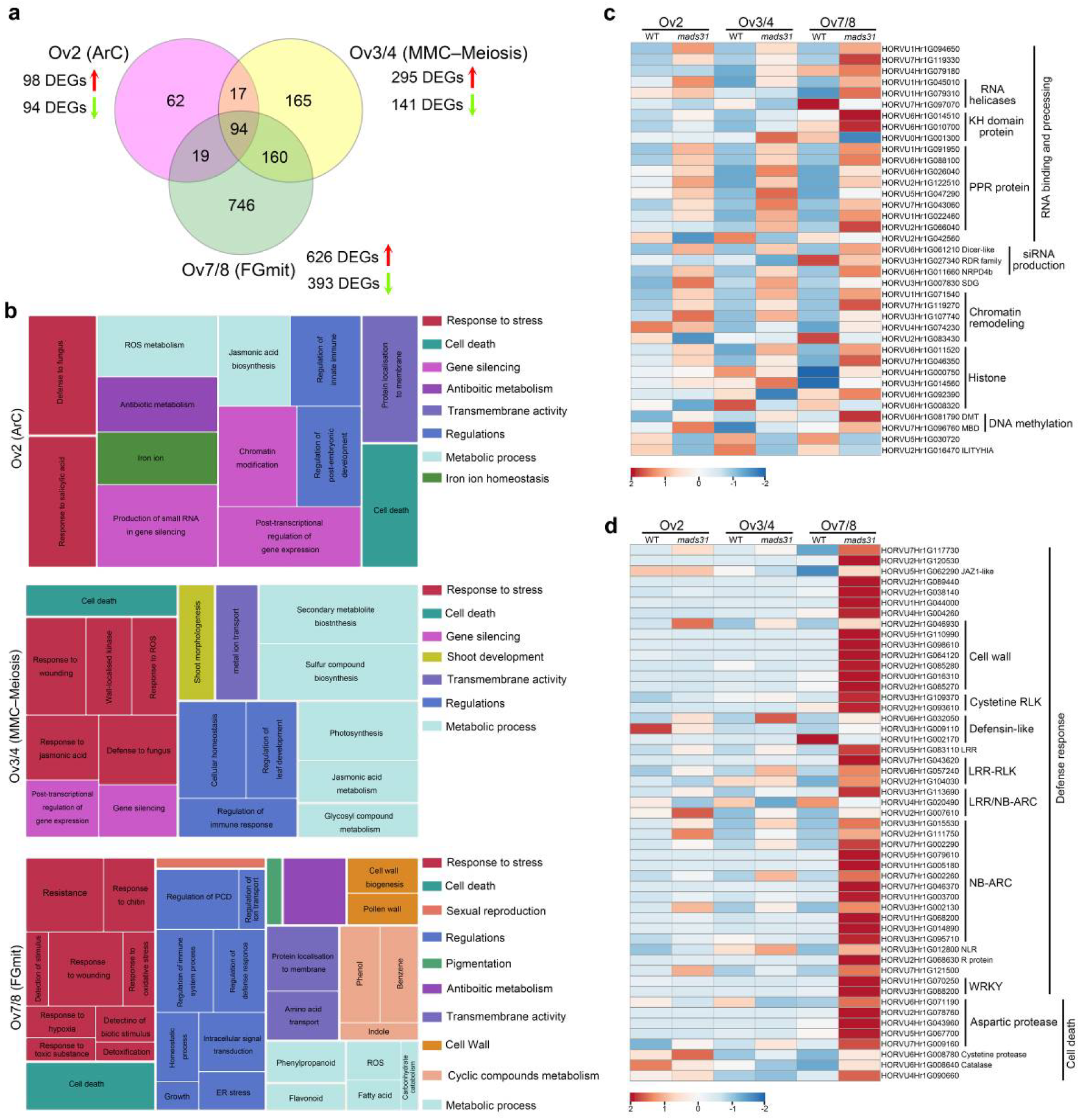
Activation of epigenetic pathways and cell death control in *mads31* ovules. **a,** Venn diagram representing the overlaps of all 1,263 DEGs identified from three developmental stages. **b,** GO enrichment of DEGs in Ov2, Ov3/4 and Ov7/8 stages. **c, d,** Heat maps of DEGs relevant to post-transcriptional and epigenetic regulation (**c**) and cell death control (**d**). KH domain, K homology domain; PPR, pentatricopeptide repeat; siRNA, small interfering RNA; RDR, RNA dependent RNA polymerase; SDG, SET domain group protein; DMT, DNA methyltransferase; MBD, methylcytosine binding domain; JAZ1, Jasmonate-ZIM; RLK, receptor-like kinase; LRR, leucine-rich repeat; NB-ARC, a functional ATPase domain; NLR, nucleotide-binding domain leucine-rich repeat containing.

The prominence of gene silencing pathway genes in the DEG lists was of particular interest, since their de-regulation has been reported to induce defects in cell proliferation and tissue development (Chen and Rechavi, 2022). DEGs encoding proteins that can bind and process RNA molecules, such as RNA helicases, K homology domain proteins, and pentatricopeptide repeat proteins, were predominantly upregulated in *mads31*, consistent with hyperactive post-transcriptional regulation. Genes encoding members of the small interfering RNA (siRNA) synthetic machinery, such as Dicer-like proteins and NPRD4b, along with factors involved in DNA methylation, histone methylation and chromatin remodeling were also upregulated (Fig. 3c and Supplementary Dataset 2). Consistent with hyperactive RNA-directed DNA methylation (RdDM) activity, a previously reported RdDM target gene *ILITYHIA* (Li et al., 2020) was down-regulated in *mads31* (Fig. 3c).

In terms of stress response and cell death pathways, upregulated genes included executors of cell death such as aspartic proteases and cysteine proteases (Fig. 3d and Supplementary Dataset 3), and a JAZ1-like gene, encoding a homologue of an inhibitor of jasmonate signaling (Chini et al., 2007). Upregulation of these genes at stage Ov7/8 correlated with increased vacuolation of the inner nucellus of *mads31* ovules, which is often a hallmark of cell death (Hara-Nishimura and Hatsugai, 2011). In addition, two *WRKY* genes, participating in both biotic and abiotic responses (Jiang et al., 2017), and genes encoding proteins involved in plant immunity, such as defensins, receptor-like kinases, NB-ARC domain proteins and cell wall remodeling proteins (Zhou and Zhang, 2020), were upregulated. Transcription factors from other families were also upregulated (Extended Data Fig. 5b), including a B3 protein and a basic helix-loop-helix (bHLH) protein, which together may indicate a cascade of transcriptional deregulation downstream of MADS31. We verified a number of these DEGs by qRT-PCR comparing wild-type versus *mads31* spikes/pistils. Consistent with cell death detected by TUNEL in later stages, we also showed that three genes encoding an aspartic protease, NB-ARC protein, and a WRKY transcription factor, were upregulated during later stages of ovule development (Ov9a and Ov9b), suggestive of prolonged defense and programmed cell death activity in *mads31* (Extended Data Fig. 5b).

In summary, loss of MADS31 appears to trigger transcriptional activation of several pathways, particularly those involved in post-transcriptional regulation, RdDM, metabolism, defense response and cell death.

### MADS31 acts to repress the post-fertilization program

To confirm whether MADS31 can function as a repressive transcription factor *in planta*, we cloned promoters from six DEGs, each predicted to carry MADS TF-binding CArG motifs and one control promoter without any CArG motif, and analyzed their activity in the presence of MADS31 via a dual-luciferase assay. Expression of MADS31 led to transcriptional repression of all six promoters, irrespective of whether the DEG was upregulated or downregulated in the *mads31* RNAseq dataset. Moreover, the number and position of the CArG motifs had minimal impact on the degree of MADS31-induced repression (Fig. 4a). Hence, MADS31 appears to be a transcriptional repressor that can act on promoters containing CArG motifs.

**Fig. 4:**
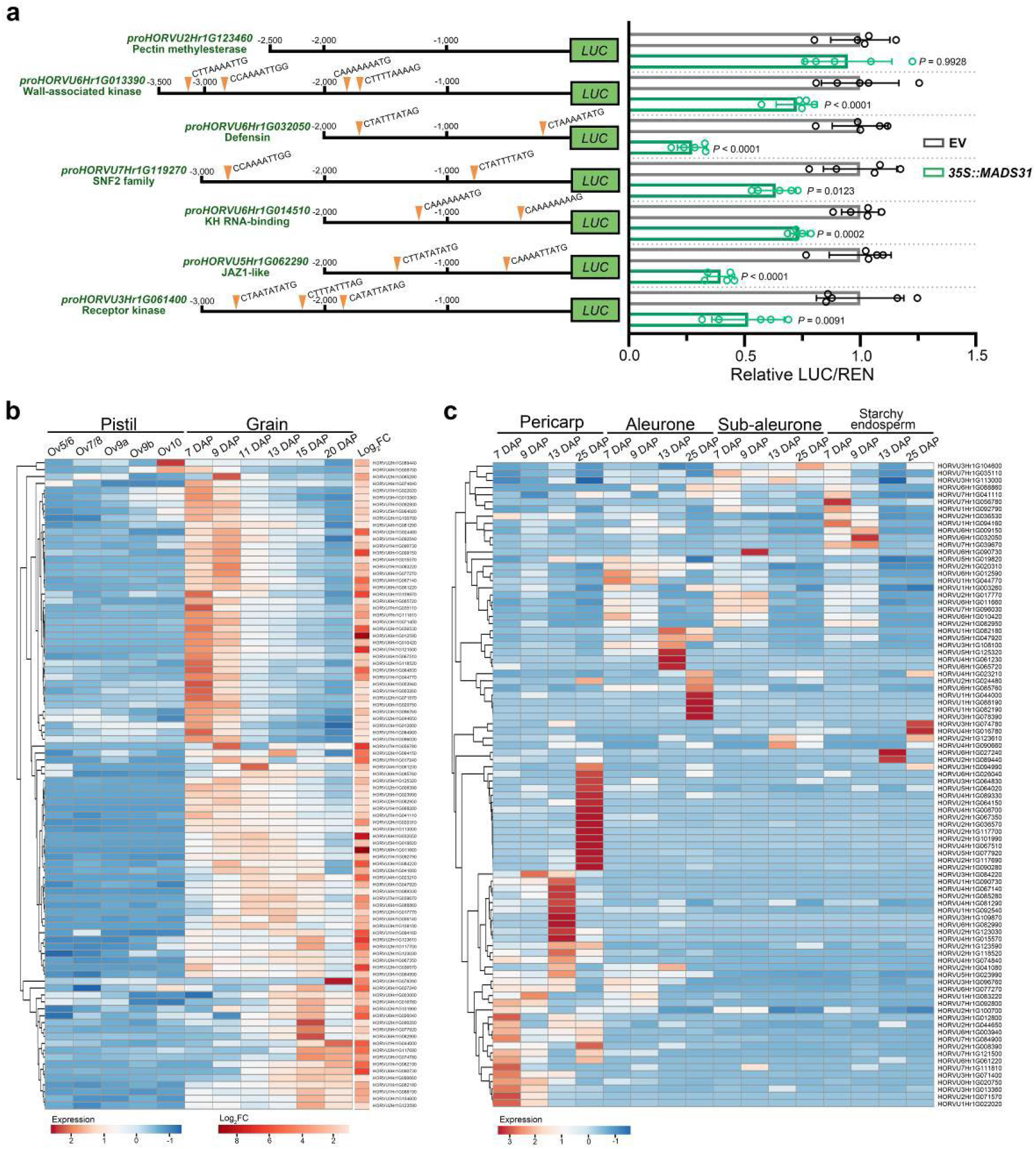
*MADS31* represses the post-fertilization program and maintains embryo sac development. **a,** Normalized luciferase activity (LUC/REN) regulated by promoters containing CArG motifs in the presence of MADS31 or empty vector (EV, negative control). The data are shown as mean ± s.d.; *n* = 5 replicates. **b,** Heat map representation of the expression patterns of 94 up-regulated DEGs in wild-type pistils and grain. Ov5/6, functional megaspore (FM) stage; DAP, days after pollination; FC, fold change. **c,** Heat map representation of the expression patterns of DEGs of **b** in wild-type grain.

Next, we examined the temporal expression profile of DEGs that are usually repressed by MADS31 (Wilkinson, 2019a; Aubert, 2018). Remarkably, of the 626 DEGs upregulated in *mads31* pistils at Ov7/8, 354 appeared to be grain-related genes predominantly expressed after fertilization during wild-type development (Supplementary Dataset 5). GO enrichment analysis of these 354 DEGs suggested a number of grain pathways are precociously activated during pre-fertilization pistil development in *mads31*, including protein modification, transmembrane transport, and phosphorus metabolism (Supplementary Dataset 5). A representative subset (94) of these genes is shown in Figure 4b, which highlights their distinct upregulation after fertilization in wild-type plants. By mapping this subset of 94 upregulated DEGs to the grain LCM transcript atlas, we identified a number of endosperm-enriched genes, including *NRPD4b* and *Defensin* (Fig. 4c). In addition, several sugar transporters and sulfotransferases that are typically expressed in the post-fertilization pericarp are activated in unfertilized *mads31* pistils. In *mads31*, these genes are expressed in the pistil at least 30 days before they would normally be up-regulated in wild-type plants (Waddington et al., 1983). These findings are consistent with MADS31 repressing transcription of a sub-class of genes involved in post-fertilization development, many of which are involved in active cell metabolism.

### Increased expression of MADS31 inhibits plant growth and modifies nucellar cell identity

The *mads31* loss of function data led us to consider the effect of increased MADS31 expression. Hence, we sought to increase MADS31 expression levels using the well-described constitutive *Ubiquitin 1* promoter of maize, which is expressed throughout the plant including in pistils and grain. Recovery of transgenic lines was severely reduced compared to other constructs, despite multiple attempts. Only two PCR positive *Ubi::MADS31* lines were identified, and these showed severe growth retardation before flowers or grains could be produced. Even after 40 days of growth post-tissue culture in soil, *Ubi::MADS31* plants exhibited excessively curled thin leaves, an absence of tillering and extreme dwarfism typified by a height of approximately 2 centimeters. After 120 days of growth, transgenic plants failed to produce any inflorescence and ultimately withered and died (Extended Data Fig. 6a). This suggests that MADS31 may act as a general repressor of growth, even beyond the ovule.

We also generated *pro::MADS31-eGFP* transgenic plants in a *mads31* and wild-type background. Importantly, *pro::MADS31-eGFP* was confirmed to be functional via complementation of the *mads31* mutant. Three *pro::MADS31-eGFP mads31/*- lines expressing MADS31-eGFP (Extended Data Fig. 7a) exhibited a wild-type phenotype in which no fertility defects or abnormal nucellus patterning were observed (Extended Data Fig. 7b–d). By contrast, extra copies of *MADS31* in a wild-type background, induced prominent changes in ovule development. An examination of mature (Ov10) ovules revealed that compared to wild-type ovules at the same stage, *pro::MADS31-eGFP* WT ovules exhibited a greater proportion of nucellar cells with characteristics typical of the inner nucellus, such as larger cell size, compressed cytoplasm, and de-esterified pectin labelling in cell walls, leading to the increased ratio of inner nucellus versus outer nucellus (Extended Data Fig. 6b–d). This suggests that increased MADS31 expression can promote "inner nucellus-like" identity.

In addition, multiple *pro::MADS31-eGFP* WT lines exhibited various degrees of dwarfism coupled with flag leaf inclination (Extended Data Fig. 6e, f), showing remarkable architectural similarities to *dicer-like 3* (Wei et al., 2014) and *osnrpd1ab* (Xu et al., 2020) mutants in rice. Previous studies of maize and Arabidopsis *argonaute* mutants have also reported that defects in the RdDM pathway can compromise nucellar cell identity (Olmedo-Monfil et al., 2010; Singh et al., 2011), reminiscent of the early defects observed in *mad31* ovules. Taken together, the *mads31* and *pro::MADS31-eGFP* data support the hypothesis that the level of MADS31 expression significantly affects ovule development, and changes in expression may interfere with the RdDM pathway.

### MADS31 controls inner nucellus development by directly repressing *NRPD4b*

Several genes involved in RdDM were identified as DEGs in *mads31* (Fig. 3c). This includes *NRPD4b*, which encodes the fourth largest subunit of RNA polymerase complex IV and V. The barley genome contains two copies of the *NRPD4* gene, *NRPD4a* (HORVU6Hr1G071930) and *NRPD4b* (HORVU6Hr1G011660). *NRPD4a* is ubiquitously expressed in pistils and grains whilst *NRPD4b* is only expressed in grains (Extended Data Fig. 8a). At the tissue level, LCM-transcript profiling revealed that *NRPD4a* transcripts accumulate in all tissues dissected from ovules and grains, but *NRPD4b* is exclusively expressed post-fertilization, mainly in the endosperm and aleurone (Extended Data Fig. 7b; Wilkinson, 2019a; Aubert, 2018). Consistent with these data, *NRPD4* expression is involved in endosperm imprinting in Arabidopsis (Klosinska et al., 2016).

*NRPD4b* transcripts were almost undetectable in wild-type barley pistils, but accumulated to high levels in *mads31* pistils of all stages (Fig. 5a). Similarly, *in situ* hybridization of *NRPD4b* detected no expression in the wild-type ovule, whilst the *mads31* ovule exhibited high expression in the nucellus, especially the basal end, with weaker expression in the ovary wall. This overlaps with the heavily reduced expression of *MADS31* in *mads31* ovules (Fig. 5b). Consistent with regulation by MADS-domain transcription factors, the promoter sequence and introns of *NRPD4b* harbor multiple CArG motifs (Fig. 5c). Using an *in vivo* assay, MADS31 protein was able to significantly repress transcriptional activity of the *NRPD4b* promoter in a dual-luciferase assay (Fig. 3d). This was also confirmed by ChIP-PCR of *pro::MADS31-eGFP* transgenic plants, which showed that MADS31 directly binds CArG motifs adjacent to the *NRPD4b* start codon (Fig. 3e).

**Fig. 5:**
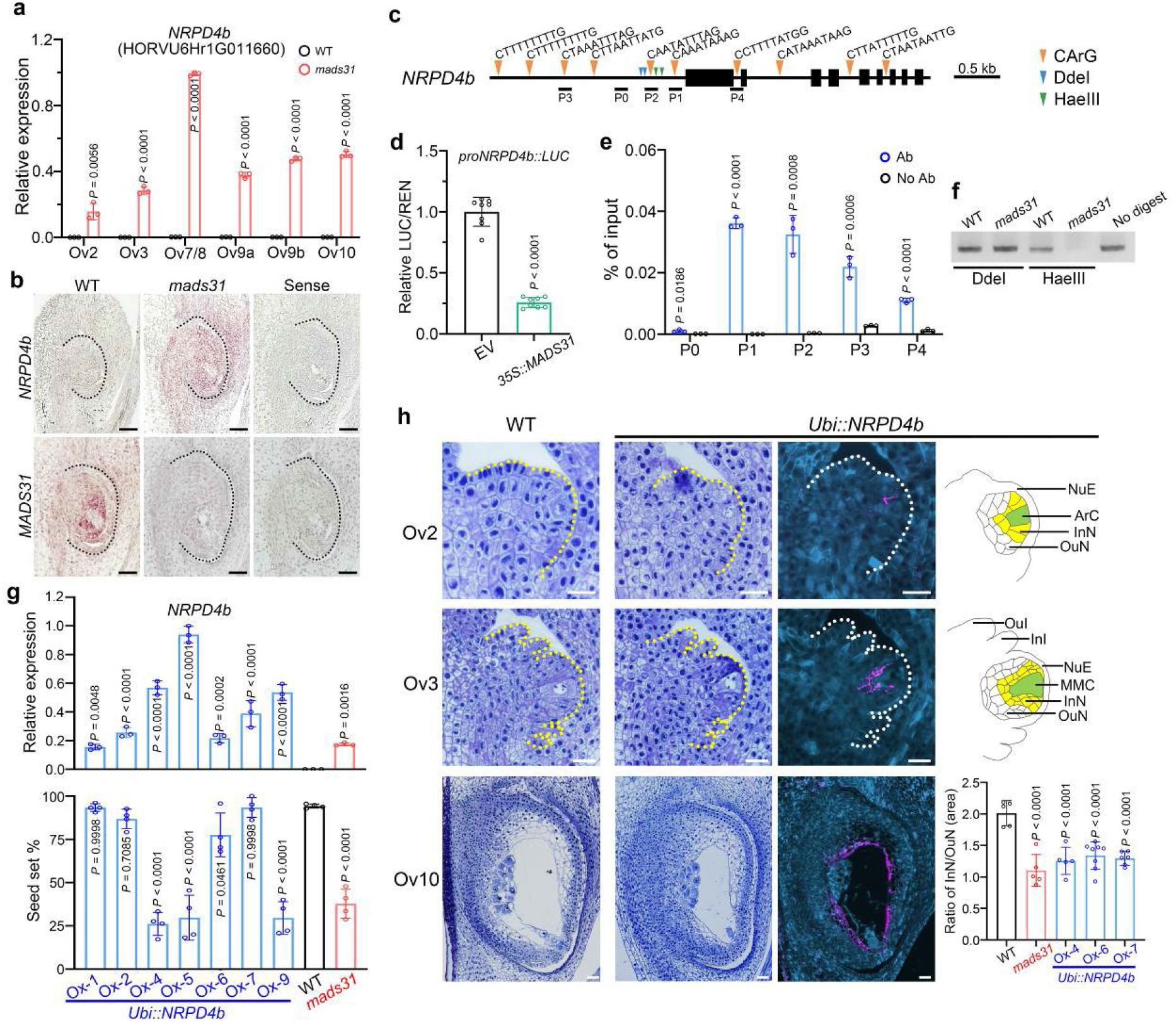
MADS31 maintains inner nucellus identity by repressing *NRPD4b* expression in the ovule. **a,** Relative expression level (RT-qPCR) of *NRPD4b* in WT and *mads31*. The data are shown as mean ± s.d.; *n* = 3 replicates. **b,** In situ hybridization of *NRPD4b* in WT and *mads31* ovules. The black dashed lines indicate ovules. Sense probe serves as negative control. Scale bars, 50 μm. **c,** The genomic region of *NRPD4b* including promoter and coding region. CArG motifs and restriction enzyme sites are marked. **d,** Normalized luciferase activity (LUC/REN) regulated by *NRPD4b* promoter in the presence of MADS31or empty vector (EV, negative control). The data are shown as mean ± s.d.; *n* = 8 replicates. **e,** Four DNA fragments with CArG motif and one without CArG motif tested by ChIP-PCR. No ab serves as negative control. The data are shown as mean ± s.d.; *n* = 3 replicates. **f,** Chop-PCR assay of DNA methylation in *NRPD4b* promoter in WT and *mads31*. Experiments were repeated three times with similar results. **g,** Overexpression of *NRPD4b* decreases seed set rate. Upper, relative expression level of *NRPD4b* in transgenic lines, wild-type and *mads31* plants. The data are shown as mean ± s.d.; *n* = 3 replicates. Lower, seeds set rates of transgenic, wild-type and *mads31* plants. The data are shown as mean ± s.d.; *n* = 4 spikes. **h,** Early (Ov2 and Ov3) and mature (Ov10) stages of wild-type and *Ubi::NRPD4b* ovules. For *Ubi::NRPD4b*, left, toluidine blue stained longitudinal sections. Middle, LM19 labeled demethylesterified pectin in the cell walls of inner nucellus. Right, diagrams of cell arrangement in the nucellus. Green, the germ cell; yellow, inner nucellus. Ratios of area of inner nucellus versus outer nucellus are shown as mean ± s.d.; *n* = 5 ovules. OuI, outer integumens; InI, inner integuments; NuE, nucellus epidermis; ArC, archesporial cell, MMC, megaspore mother cell; InN, inner nucellus, OuN, outer nucellus. Scale bars, 25 μm.

We considered how the repression of *NRPD4b* by MADS31 might be achieved. Previous studies have described that the methylation status of promoter regions influences the transcription of adjoining sequences (Zhang et al., 2018). Several restriction nuclease cut sites, including *DdeI* and *HaeIII*, are sensitive to cytosine methylation (Li et al., 2020) and were identified flanking the CArG motifs in the *NRPD4b* promoter. We performed chop-PCR and identified a loss of methylation in *HaeIII* sites close to one of the CArG motifs in *mads31* (Fig. 5f), which was consistent with upregulation of *NRPD4b*. To uncouple this regulation of the *NRPD4b* promoter by MADS31, we created transgenic *Ubi::NRPD4b* plants that overexpress *NRPD4b* in a wild-type background. The resulting lines exhibited lower seed set similar to that observed in *mads31* (Fig. 5g). Moreover, in lines highly expressing *NRPD4b*, ovules showed similarities to those from *mads31* plants, as typified by inner nucellar cells being irregular in shape, disorganised, as well as containing less de-esterified pectin in their cell walls at stages Ov2 and Ov3 (Fig. 5h and Extended Fig. 9a). At maturity (Ov10), *Ubi::NRPD4* transgenic ovules showed altered patterning with the increased ratio of inner nucellus versus outer nucellus, compared to wild-type, with less nucellar cells labelled by LM19, albeit more than *mads31* (Fig. 5h and Extended Fig. 9b). Thus, *Ubi::NRPD4b* plants show phenotypes similar to those of a weak *mads31* phenotype (Fig. 5h).

The RdDM pathway is known to recruit histone methyltransferases for the methylation of H3 on the lysine 9 residue (H3K9), thereby reinforcing a repressive transcriptional state (Du et al., 2014; Li et al., 2018). Immunodetection of histone modifications is routinely used to investigate the consequences of RdDM within ovules (She et al., 2013; Oliver et al., 2013) (Fig. 6a). Consistent with the increased transcription of *NRPD4b* in both *mads31* and *Ubi::NRPD4b* ovules, nucellar cells surrounding the embryo sac (Fig. 6b) exhibited a greater degree of repressive H3K9me2 labelling compared to wild-type (Fig. 6d). We also investigated differences in H3K27me1 deposition, which is believed to occur independent of DNA methylation (Mathieu et al., 2005). At ovule stage Ov7/8, wild-type ovules exhibited an even distribution of H3K27me1 immunolabelling throughout the nucellus. At the same stage in *mads31* and *Ubi::NRPD4b* ovules, inner nucellar cells surrounding the germline showed significantly reduced H3K27me1 labelling (Fig. 6c, e). Taken together, increased H3K9me2 labelling suggests that *NRPD4b* overexpression affects the global methylation pattern of the inner nucellus cells. Intriguingly, the altered H3K27me1 immunolabelling in the same region is the inverse of H3K9me2, and may reflect downstream changes in cell identity. It is also consistent with the independent regulation of H3K9me2 and H3K27me1 (Liu et al., 2010).

**Fig. 6:**
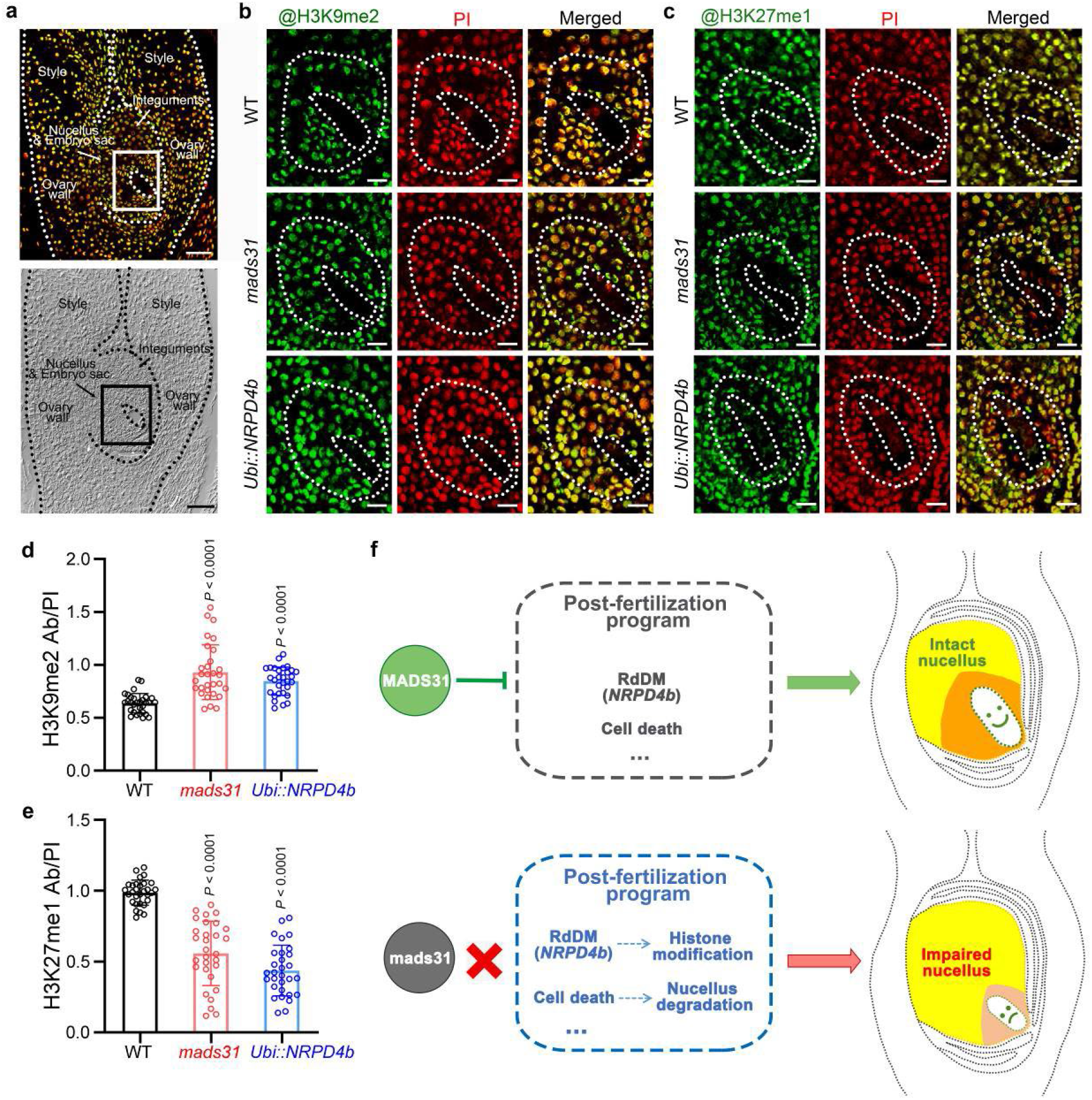
Upregulation of *NRPD4b* alters the distribution of histone marks. **a,** The whole pistil labeled by antibody to detect histone modification, in the fluorescent channel (upper) and DIC channel (lower). The white and black rectangles indicate the region of interest shown in **b** and **c**. Scale bar, 50 μm. **b,** Immunolabeling of H3K9me2 in wild-type, *mads31* and *Ubi::NRPD4b* ovules at Ov7/8 stage. Scale bars, 25 μm. **c,** Immunolabeling of H3K27me1 in wild-type, *mads31* and *Ubi::NRPD4b* ovules atOv7/8) stage. White dash lines indicate the embryo sac and inner nucellus regions. PI, propidium iodide. Scale bars, 25 μm. **d,** Relative H3K9me2 modification level (measured as antibody signal intensity/DNA signal intensity) in the inner nucellus region of wild-type, *mads31* and *Ubi::NRPD4b* ovules. The data are shown as mean±s.d.; *n* = 30 nuclei. **e,** Relative H3K27me1 modification level (measured as antibody signal intensity/DNA signal intensity) in the inner nucellus region of wild-type, *mads31* and *Ubi::NRPD4b* ovules. The data are shown as mean ±s.d.; *n* = 30 nuclei. **f,** Proposed model of MADS31 nucellus patterning. In wild type, MADS31 is preferentially expressed in the inner nucellus, repressing post-fertilization activities such as gene silencing and nucellus degradation to maintain the tissue integrity and support embryo sac development. In *mads31* ovules, the lack of repression from MADS31 causes activation of *NRPD4b* and cell death control genes, which further alters histone modifications and accelerates tissue degeneration, respectively.

## Discussion

Female germline cells in the plant ovule are enveloped in layers of somatic tissue, providing multiple sources of regulatory cues for germline development. These include autonomous pathways acting only in the germline, and distal non-cell autonomous pathways acting from the nucellus, integuments, funiculus, or vascular system. In terms of local control, the nucellus represents the closest source of somatic information to coordinate and/or transduce information for germline progression. In Arabidopsis, the nucellus is prominent during early ovule growth, but degrades quickly and occupies only a small proportion of the ovule after meiosis (Lu and Enrico, 2018), leaving room for the integuments to directly enclose the embryo sac. By contrast, cereal ovules exhibit a larger multiple-layered nucellus, and the germline is surrounded by a prominent nucellus until fertilization, meaning the integuments have no direct connection to the embryo sac. Here, we show that the nucellus of barley can be further differentiated into two subtypes, the inner nucellus and the outer nucellus, based on their position relative to the germline, their cellular morphology, and cell wall organization. MADS31 is a Bsis MADS-box protein that is preferentially expressed in the inner nucellus. The loss of MADS31 leads to hyperactive RdDM activity mediated by *NRPD4b* and premature cell death, impairing the integrity of nucellus and germline development (Fig. 6d). Thus, MADS31 represents a key factor to dissect the mechanistic role of different nucellus tissues in cereal ovule and seed development.

A differentiated “inner zone” within the nucellus is not unique to Triticeae cereals, and has been reported in other angiosperms such as soybean (Folsom and Cass, 1988) and Ginkgo (Douglas et al., 2007), sometimes referred to as a nucellar epithelium. The bond between the inner nucellus and female germline shares noteworthy similarities with that of the tapetum and male germline in anthers. Both the inner nucellus and tapetum provide a transient interface between sporophytic and germline tissues (Rudall, 2021), and both undergo precisely controlled cell death, which is required for downstream development (Russell, 1979; Nogueira et al., 2021, Parish and Li, 2010; Radchuk et al., 2006). MADS31 protein accumulates in the nucellus but not the germline (Fig. 1c), and defective germline formation in *mads31* mutants correlates with altered inner nucellus development. Hence, MADS31 appears to maintain inner nucellus identity, which is essential for female germline development.

The molecular interplay between the germline and surrounding cells during sporogenesis involves complex regulatory activities (Pinto et al., 2019; Marchant and Walbot, 2022). Recent findings in the anther suggest that so-called nurse cells in the tapetum provide siRNAs for RdDM activity in the male germline (Long et al., 2021). In the Arabidopsis ovule, RdDM pathways are thought to generate siRNAs in the nucellar cells to coordinate MMC specification (Olmedo-Monfil et al., 2010; Singh et al., 2011; Rodríguez-Leal et al., 2015; Su et al., 2017; Su et al., 2020). Despite this, remarkably little is known about the upstream regulators of tissue-specific RdDM activities. In Arabidopsis, the D-class MADS-box gene *STK* activates the biogenesis of siRNAs via RDR6 and AGO9 to restrict the expression of SPOROCYTELESS/NOZZLE (SPL/NZZ), leading to the specification of a single female germline cell (Mendes et al., 2020). Here we show that MADS31 is potentially a negative regulator of the RdDM pathway, and acts by directly repressing *NRPD4b* expression. Increased *NPRD4b* expression in *mads31* and *Ubi::NRPD4b* ovules interrupts inner nucellus development, leading to altered H3 methylation patterns (Fig. 5 and 6). Thus, MADS31 appears to directly regulate RdDM related pathways required for germline development. In light of STK D-class MADS function in promoting expression of RdDM components, this suggests that sophisticated regulation of the RdDM pathway by different classes of MADS-box genes may be important to maintain a balance between somatic and germline identity.

Bsis MADS-box genes exist widely in plants that bear female reproductive organs (Becker et al., 2002, Yang et al., 2016; Hu et al., 2021), but show relatively low cross-species homology at the protein level. Despite this, Bsis class members are typically expressed in somatic cells of ovules, such as the nucellus, integuments/ovule envelopes and carpel/ovary wall. Due to the lack of mutant resources in many species, functional research of Bsis class has been limited to model plants. In Arabidopsis, both Bsis genes, *GOA* and *ABS*, are expressed in nucellus and integuments, although integument development is more sensitive to changes in their function (Nesi et al., 2002; Prasadet al., 2010). In rice and barley, *MADS29* is also expressed in the nucellus and integuments, but the loss of *MADS29* predominantly results in changes to the nucellus and nucellar projection (Lee et al., 2013, Shoesmith et al., 2021). Here, we show that MADS31 regulates inner nucellus development, also with limited impact on the integuments. We speculate that as ovule development progresses, Bsis genes primarily regulate the development of tissue adjoining the germline, which is the integuments in tenuinucellar ovules and the nucellus in crassinucellar ovules. Although detailed molecular mechanisms explaining Bsis function are mostly unknown, Bsis homologs exhibit an interesting common trait in ectopic expression experiments. Overexpression of *GOA*, *ABS*, *PeMADS28* and *GbMADS9* by a constitutive promoter in Arabidopsis leads to phenotypes such as dwarfism, abnormal floral organs, early flowering time and sterility (De Folter et al., 2006; Erdmann et al., 2010, Yang et al., 2016; Shen et al., 2021). Similarly, transgenic plants overexpressing *OsMADS29* in rice are dwarfed, early flowering and sterile (Nayar et al., 2013). Here, the overexpression of *MADS31* in barley has extreme effects on development, whereby transgenic plants are dramatically stunted and unable to flower. Thus, Bsis class members appear to be negative regulators of growth. Their restricted expression in the ovule may therefore limit growth in key zones adjoining the germline niche and in nutrient transfer tissues, creating an optimal environment for germline initiation and nutritional support.

Consistent with this hypothesis, we determined that MADS31 predominantly represses expression of target genes (Fig. 4). Repressive transcription factors are an essential component of growth and development, often interacting with chromatin remodeling proteins to stabilize genomic structure and enable correct timing of target gene expression. The molecular consequences of de-repression in *mads31* are remarkable, and include precocious activation of cell death pathways and seed-specific genes such as the endosperm-specific RdDM component *NRPD4b* in unfertilized ovules. This de-repression coincides with premature cell death in the inner nucellus and potential rewiring of metabolic pathways. During normal ovule development, fertilization is an important cue for ovules to degrade maternal tissues, prior to initiating seed development. The altered nucellus morphology in *mads31* may therefore result from the precocious activation of programmed cell death pathways. Whether this is the direct cause of germline abortion is unclear; a non-mutually exclusive possibility is that premature expression of *NRPD4b* compromises RNA polymerase complex IV/V function and siRNA pathways, thereby interfering with the function of the inner nucellus as a nurse tissue for the female germline.

Taken together, our findings suggest MADS31 is required to maintain timely transitions during nucellus growth and differentiation. MADS-box proteins typically form heteromeric complexes to execute function (Ng and Yanofsky, 2001) and a physical interaction between OsMADS29 and OsMADS31 suggests the Bsis class proteins can function in a complex (Nayar et al., 2014). Although MADS29 targets remain unknown in barley, rice OsMADS29 has been reported to activate NBS-LRR and cysteine proteases involved in stress response and cell degeneration (Yin and Xue, 2012; Nayar et al., 2013). Here we show that similar genes appear to be repressed by MADS31, suggesting that MADS31 and MADS29 may act antagonistically in controlling cell death. How MADS31 establishes a repressive state remains unclear, although MADS-box proteins have been shown to interact with chromatin remodelers to regulate their targets (Masiero et al., 2002; Abraham-Juárez et al., 2020; Lai et al., 2021). The decrease of DNA methylation in the *NRPD4b* promoter region suggest that this RdDM component may itself be a target of the epigenetic silencing machinery. However, unlike LEAFY, which acts as a master activator (Lai et al., 2021) by making chromatin more accessible, MADS31 would hypothetically interact with epigenetic components to reinforce repressive states.

## Methods

### Plant materials and creation of transgenic plants

The wild type barley (*Hordeum vulgare*) cultivar Golden Promise (GP) was used as a control and donor plant for transgenesis. An optimized CRISPR/Cas9 genome editing system was utilized to generate mutants (Ma et al., 2015). Two specific targets were selected for *MADS31* near the start codon. Targets were sequenced in GP to guarantee paring between single guide RNA (sgRNA) and genomic DNA. SgRNA–target 1 (T1) was driven by rice promoter *OsU6c*, and sgRNA–T2 was driven by rice promoter *OsU3*. The sgRNA expression cassettes of *OsU6c–sgRNA–T1* and *OsU3–sgRNA–T2* were amplified from pYLsgRNA–OsU6c and pYLsgRNA–OsU3 plasmids using Phusion High-Fidelity DNA Polymerase (New England BioLabs) and cloned into a binary vector, pYLCRISPR–Cas9Pubi–H, using two *BsaI* sites as described.

To trace MADS31 protein accumulation in planta, a ~4kb genomic DNA fragment including the promoter and full genomic coding region of *MADS31* were fused in frame to *eGFP* and inserted between the *HindIII* and *BstEII* sites of pCAMBIA1301, using In-Fusion (Takara) cloning technology. This construct was also used to complement the *mads31* mutant. For *MADS31* overexpression, the full length *MADS31* coding sequences was inserted into vector pU1301 via *KpnI* and *BamHI* sites behind the maize *Ubiquitin 1* promoter. The same cloning method was used for *NRPD4b* overexpression. All primers used for constructs are listed in Supplemental Table 1.

All constructs were transformed into immature GP or *mads31* embryos using an *A. tumefaciens* AGL1-mediated transformation method described previously (Bartlett, 2008). All barley plants were grown in cocopeat soil, in 15 °C light,12 °C dark, 16 hours daylight with 70% humidity (The Plant Accelerator, Waite Campus, The University of Adelaide, Australia). Individual T0 plants carrying homozygous or biallelic mutations generated by CRISPR were identified by Sanger sequencing (AGRF) of the targets and flanking region amplified by the Phire Plant Direct PCR Kit (ThermoFisher Scientific).

### Plant and pistil phenotyping

Barley plants, spikes, anthers and pistils were photographed using a Nikon D5600 digital camera. Mature anthers were dissected from spikelets and crushed on microscopy slides to release pollen grains. Pollen grains were stained in Lugol's iodine for 30 seconds and photographed using an optical microscope (Ni-E, Nikon). For clearing, whole pistils were collected from spikelets approaching anthesis and fixed in ice-cold FAA immediately. Pistils were dehydrated in a series of 70, 80, 90, and 100% (v/v) ethanol and cleared in Hoyer's solution for four weeks as previously described (Wilkinson, 2017). Cleared pistils were imaged using a Zeiss AxioImager M2 with differential contrast microscopy (DIC).

Fresh spikes (for ovules of Ov2–Ov7/8) or pistils (for ovules of Ov9b/10) were collected from *pro::MADS31-eGFP* plants and embedded in 5% (m/v) agarose blocks immediately. After solidifying and trimming, samples were sectioned into 50-70 μm slices using a Leica Vibratome VT1200. Slices were laid on microscopy slides and mounted by 50% (v/v) glycerol solution. Ovule sections containing the germline were imaged using an A1R Laser Scanning Confocal Microscope (Nikon) (eGFP, excitation, 488 nm; emission, 505-520 nm). Images were processed with NIS-Elements Viewer 4.20 (Nikon). Intact ovules of Ov3–Ov10 were carefully dissected from pistils of *pro::MADS31-eGFP* and *mads31/pro::MADS31-eGFP* plants and photographed with a Zeiss AxioImager M2 (eGFP, excitation, 450-490 nm; emission, 500-550 nm; auto fluorescence, excitation, 335-383 nm; emission 420-470 nm).

### Sectioning and pectin immunolabeling

Pistils or whole spikelets from wild-type plants, mutants and transgenic plants were collected and fixed in FAA, dehydrated in a series of 70, 80, 90, and 100% (v/v) ethanol and embedded in Technovit 7100 resin (Kulzer Technique) as described by the manufacturer. Samples were sectioned to a thickness of 1.5 μm on a Leica Ultramicrotome. Sections were stained in 0.5% Toluidine Blue (w/v) and imaged by Nikon Ni-E optical microscope. For pectin immunolabeling, unstained sections were incubated with rat antibody LM19 (1:100 dilution, PlantProbes), followed by secondary antibody Alexa Fluor 555 conjugated goat anti-rat IgG (1:200 dilution, Invitrogen) as described previously (Betts et al., 2017). Then sections were stained in Calcofluor White Stain (Sigma-Aldrich) for 1 min for background cell wall labeling. After three rinses with water, sections were mounted with 90% glycerol and imaged by a Zeiss AxioImager M2 (LM19, excitation, 538–562 nm; emission, 570–640 nm; calcofluor stain, excitation, 335–383 nm; emission 420–470 nm). Fluorescence intensity was measured using ImageJ. Mean fluorescence of background was measured to calculate corrected total cell fluorescence (CTCF).

### TUNEL assay

Tissues were harvested into glass vials and fixed in ice cold FAA. Plant materials were dehydrated in a series of 70, 80, 90, and 100% (v/v) ethanol and embedded in paraffin as described previously (Wu, 2012). 8 μm paraffin sections were prepared using a Leica rotary microtome RM2265 and transferred to polysine coated slides (ThermoFisher Scientific), dewaxed, rehydrated and post-fixed in 4% (w/v) paraformaldehyde. Nick-end labeling of nuclear DNA fragmentation mediated by Terminal Deoxynucleotidyl Transferase (TdT) was performed following the instructions of DeadEnd^™^ Fluorometric TUNEL System (Promega). Sections were stained in 1 μg/ml propidium iodide (PI) before mounted by 90% (v/v) glycerol with 25 mg/ml DABCO, then imaged by an A1R Laser Scanning Confocal Microscope (Nikon) (fluorescein-12-dUTP, excitation, 395 nm; emission, 500–540 nm; PI, excitation, 561 nm; emission, 590–640 nm).

### RNA extraction and RT-qPCR

Total RNA was extracted from barley tissues using a Spectrum^™^ Plant Total RNA Kit (Sigma-Aldrich). 2 μg of total RNA was purified using the TURBO DNA-*fee*^™^ Kit to remove genomic DNA. First-strand cDNA was generated using SuperScript IV Reverse Transcriptase (Invitrogen) and oligo-dT primer, following the manufacturer's instructions. Diluted cDNA was used as templates mixed with iTaq Universal SYBR Green Supermix (Bio-Rad) for real-time quantitative PCR using a QuantStudio Flex 6 (Life Technologies) machine. *HvACTIN7* was used as housekeeping gene for normalization. All primers used for qRT-PCR are listed in Supplemental Table 1.

### RNA in situ hybridization

*MADS31* and *NRPD4b*-specific fragments were amplified from cDNA templates by PCR using primers fused with T7 polymerase promoters. PCR products were used as DNA templates for in vitro transcription. Digoxigenin (DIG)-labeled NTP (Roche) was used to label antisense and sense probes generated by T7 polymerase (ThermoFisher Scientific), according to the manufacturer's instructions. Barley tissues were fixed, embedded and sectioned as described in TUNEL assay. Rehydration, post-fix, hybridization, stringent washes and immunodetection were automatically performed in an InsituPro VSi robot (Intavis). To visualize hybridization signal, an antibody conjugate anti-DIG-AP (1:1000 dilution, Roche) and NBT/BCIP substrate were used for coloring reaction. All primers used are listed in Supplemental Table 1.

### RNA-sequencing and data analysis

In wild type and *mads31*, for samples of Ov2 and Ov3/4 stages, whole barley spikes at Waddington scale (Waddington et al., 1983) W5.5–6 and W6.5–7 stages were collected, respectively. Anthers were carefully removed by a dissecting needle. For samples of Ov7/8 stages, pistils at W8.5–8.75 stage were dissected from spikelets. Total RNA was isolated from tissues described above for each of three biological replicates using a RNeasy Plant Mini Kit (Qiagen). RNA quality assessment, libraries preparation and paired-end sequencing were performed at Novogene (Australia). The analysis pipeline of RNA-seq data was exerted following the description by Li et al. (Li et al., 2021). After trimming adaptors and filtering, clean reads were mapped to the barley reference genome (Mascher et al., 2017) using HISAT2 aligner. FPKM (fragments per kilobase per million) were normalized by HTSeq. Genes were considered as differentially expressed at a false-discovery rate-adjusted *P* value < 0.05 and a log2FoldChange >1 or < −1. A total of 1,263 DEGs were identified by the R package DESeq2 and further annotated according to BLASTX against protein databases of Arabidopsis (https://www.arabidopsis.org) and rice (http://rice.uga.edu). Venn diagram was created based on DEGs identified. GO analysis was performed according to Li et al., 2021. The R package ClusterProfiler was utilized for GO enrichment and REVIGO (Supek et al., 2011) was used to build treemaps for visualizing enriched GO terms. For gene expression heatmaps, the original FPKM or TPM values of genes of interest were extracted from RNA-seq data or the LCM-RNA-seq data (Wilkinson, 2019a; Aubert, 2018). The expression heatmap was created using ClustVis (Metsalu et al., 2015).

### ChIP-PCR

1 g of material including young spikelets (W7.5-8.5) with anthers pinched off, and pistils (W8.75-10) were harvested from *pro::MADS31-eGFP* transgenic plants. ChIP was performed following the method previously described by Li et al., 2021. In brief, chromatin was cross-linked, isolated by nuclei lysis and sonicated into ~100-500bp, centering around 250bp. Sheared chromatin was pre-cleared by salmon sperm (SS) DNA/Protein A/G agarose beads (Thermo Fisher Scientific) before overnight incubation with anti-GFP antibody (ABclonal) at 4 °C. The Protein A/G agarose beads were added for 2 h incubation, then washed in low salt, high salt, LiCl and TE buffer. The beads were washed twice by elution buffer to collect immunocomplexes. Reverse crosslinking was performed in 0.2M NaCl solution at 65 °C for overnight incubation. DNA was purified by proteinase K digestion, phenol-chloroform-isoamyl alcohol extraction and precipitation with ethanol, sodium acetate (pH 5.2), and glycogen at - 80 °C. Purified input and immunoprecipitated DNA were used as templates for qPCR to calculate enrichment. "No antibody" precipitation was used as negative control. All primers are listed in Supplementary Table 1.

### Dual-luciferase assay

The dual-luciferase assay was performed using transient expression in *N. benthamiana* leaves, following a method described previously (Li et al, 2014). Effector plasmid was constructed by inserting the full length *MADS31* coding region into the *HindIII* and *BamHI* sites of the pGreenII-0000 vector which drives effector expression by 35S promoter. A series of promoters of DEGs were amplified from genomic DNA and cloned into the *HindIII* and *BamHI* sites of pGreenII-0800-LUC vector to drive expression of the *LUC* reporter gene. All plasmids including empty vector pGreenII-0000 were co-transformed with helper plasmid pSoup-P19 into *A. tumefaciens* GV3101 cells. Full-strength overnight Agrobacterium cultures were collected and resuspended. Each reporter strain was mixed with MADS31 effector strain or empty vector strain at a ratio of 1:4 (v:v), respectively. The reporter-effector mixture was infiltrated into young tobacco leaves using a 2 ml syringe, then plants were kept in weak light for 48 h. Leaves were harvested and processed using the Dual-Luciferase^®^ Reporter Assay System (Promega), following the manufacturer's instructions. *Renilla* luciferase was used as an internal control to normalize firefly luciferase. All primers are listed in Supplementary Table 1.

### Chop-PCR

For Chop–PCR (DNA methylation-sensitive restriction endonuclease digestion followed by PCR), barley genomic DNA was extracted from wild-type and *mads31* pistils (W8–8.75), using the CTAB (cetyltrimethylammonium bromide) method (Dasgupta et al., 2019). 1 μg genomic DNA was digested overnight by DNA methylation-sensitive restriction endonuclease DdeI and HaeIII (New England BioLabs) and used as a template for PCR reactions using primers flanking the endonuclease recognition sites. DdeI and HaeIII report on CHH methylation, where H indicates A, T, or C. Non-digested genomic DNA was served as control. All primers are listed in Supplementary Table 1.

### Histone immunolabeling

Immunodetection of histone methylation was performed following a previous method with modifications (Nic-can et al., 2013). Tissues were fixed, embedded and sectioned as described in TUNEL assay. After dewaxing and rehydration, paraffin sections (6 μm) were microwave-heated in 10 mM citrate buffer (pH 6.0) for 5 min at high power for antigen retrieval. Sections were incubated with blocking buffer [3% (m/v) BSA in PBS buffer] for 1 h before overnight incubation with primary antibodies for H3K9me2 (1:400 dilution, abcam) and H3K27me1 (1:600 dilution, ThermoFisher Scientific) at 4°C in a humidified chamber. Alexa Fluor 488 conjugated anti-mouse and anti-rabbit IgG (1:400 dilution, Invitrogen) were used as secondary antibodies to visualize immunosignals. Sections were counterstained in 1 μg/ml PI, rinsed in water and imaged by an A1R Laser Scanning Confocal Microscope (Nikon) (AF488, excitation, 488 nm; emission, 505–520 nm; PI, excitation, 561 nm; emission, 590–640 nm). The primary antibodies were omitted as the negative control. Antibodies signals and PI signals were measured using ImageJ.

### Statistical analysis and replication

For all cytological analysis in ovules, including measurement of embryo sac area, resin sections and pectin immunolabelling, 50–100 spikelets (for observation of young ovules) or pistils (for observation of mature ovules) were collected from 4–6 barley plants of various genotypes and fixed in FAA. A certain number of spikelets or pistils were randomly picked for various experiments; the exact numbers of ovules used for cellular morphology and fluorescence intensity measurement are shown in figures or figure legends. If not specified, at least 3 ovules were observed, and representative images was shown in figures.

For experiments using paraffin sections, including in situ hybridization, TUNEL assay and histone methylation immunolabelling, 50–100 spikelets or pistils were collected from 4–6 replicate barley plants of each genotype and fixed in FAA. At least 20 spikelets or pistils for each genotype were randomly picked and embedded in paraffin. Experiments were performed three times using >3 samples for each repeat. All technical replicates showed similar results, and representative images were shown in figures. For RNA extraction and ChIP experiments, 200-1000mg spikelets or pistils were collected from 4-6 barley plants of various genotypes, except that samples were collected from each plant of *NRPD4b* overexpression lines. For qRT-PCR and ChIP-PCR, at least three technical replicates were performed.

For Dual-LUC, a whole tobacco leaf was infiltrated by *A. tumefaciens* culture. A puncher with 5 mm diameter was used to harvest 5-8 pieces of samples as replicates for luciferase activity assay.

For eGFP signal observation in transgenic lines, at least three ovules were examined from each of three plants. Representative images were shown in figures.

For seed set percentages, 4-9 barley plants of various genotype were used as biological replicates, as indicated in figures or figure legends. In the case of *NRPD4b* overexpression lines, 4 spikes from each plant were used as biological replicates.

GraphPad Prism 9 was used for generating graphs and statistical analysis. Statistical methods were described in figure legends and exact *P* values were shown in figures.

## References

Abraham-Juárez, M.J., Schrager-Lavelle, A., Man, J., Whipple, C., Handakumbura, P., Babbitt, C. and Bartlett, M., 2020. Evolutionary variation in MADS box dimerization affects floral development and protein abundance in maize. Plant Cell, 32(11), pp.3408–3424.

Aubert, M.K., 2018. Molecular and genetic characterisation of early aleurone development in barley (Hordeum vulgare L.) (Doctoral dissertation).

Becker, A., Kaufmann, K., Freialdenhoven, A., Vincent, C., Li, M.A., Saedler, H. and Theissen, G., 2002. A novel MADS-box gene subfamily with a sister-group relationship to class B floral homeotic genes. Molecular Genetics and Genomics, 266(6), pp.942–950.

Bowman, J.L., Mansfield, S.G., Modrusan, Z., Reiser, L., Fischer, R.L., Haughn, G.W., Feldman, K.A. and Webb, M.C., 1994. Ovules. In Arabidopsis (pp. 297–331). Springer, New York, NY.

Brambilla, V., Battaglia, R., Colombo, M., Masiero, S., Bencivenga, S., Kater, M.M. and Colombo, L., 2007. Genetic and molecular interactions between BELL1 and MADS box factors support ovule development in Arabidopsis. The Plant Cell, 19(8), pp.2544–2556.

Ceccato, L., Masiero, S., Sinha Roy, D., Bencivenga, S., Roig-Villanova, I., Ditengou, F.A., Palme, K., Simon, R. and Colombo, L., 2013. Maternal control of PIN1 is required for female gametophyte development in Arabidopsis. PloS one, 8(6), p.e66148.

Chen, X. and Rechavi, O., 2022. Plant and animal small RNA communications between cells and organisms. Nature Reviews Molecular Cell Biology, 23(3), pp.185–203.

Chini, A., Fonseca, S.G.D.C., Fernandez, G., Adie, B., Chico, J.M., Lorenzo, O., García-Casado, G., López-Vidriero, I., Lozano, F.M., Ponce, M.R. and Micol, J.L., 2007. The JAZ family of repressors is the missing link in jasmonate signalling. Nature, 448(7154), pp.666–671.

Daneva, A., Gao, Z., Van Durme, M. and Nowack, M.K., 2016. Functions and regulation of programmed cell death in plant development. Annual Review of Cell and Developmental Biology, 32, pp.441–468.

Dasgupta, P. and Chaudhuri, S., 2019. Analysis of DNA methylation profile in plants by Chop-PCR. In Plant Innate Immunity (pp. 79–90). Humana, New York, NY.

De Folter, S., Shchennikova, A.V., Franken, J., Busscher, M., Baskar, R., Grossniklaus, U., Angenent, G.C. and Immink, R.G., 2006. A Bsister MADS - box gene involved in ovule and seed development in petunia and Arabidopsis. The Plant Journal, 47(6), pp.934–946.

Douglas, Andrew W., Dennis W. Stevenson, and Damon P. Little. “Ovule development in Ginkgo biloba L., with emphasis on the collar and nucellus.” International Journal of Plant Sciences 168.9 (2007): 1207–1236.

Du, J., Johnson, L.M., Groth, M., Feng, S., Hale, C.J., Li, S., Vashisht, A.A., Gallego-Bartolome, J., Wohlschlegel, J.A., Patel, D.J. and Jacobsen, S.E., 2014. Mechanism of DNA methylation-directed histone methylation by KRYPTONITE. Molecular cell, 55(3), pp.495–504.

Endress, P.K., 2011. Angiosperm ovules: diversity, development, evolution. Annals of Botany, 107(9), pp.1465–1489.

Erdmann, R., Gramzow, L., Melzer, R., Theißen, G. and Becker, A., 2010. GORDITA (AGL63) is a young paralog of the Arabidopsis thaliana Bsister MADS box gene ABS (TT16) that has undergone neofunctionalization. The Plant Journal, 63(6), pp.914–924.

Folsom, M.W. and Cass, D.D., 1988. The characteristics and fate of the soybean inner nucellus. Acta botanica neerlandica, 37(3), pp.387–393.

Frohlich, M.W., 2003. An evolutionary scenario for the origin of flowers. Nature Reviews Genetics, 4(7), pp.559–566.

Hara-Nishimura, I. and Hatsugai, N., 2011. The role of vacuole in plant cell death. Cell Death & Differentiation, 18(8), pp.1298–1304.

Hu, J., Chang, X., Zhang, Y., Yu, X., Qin, Y., Sun, Y. and Zhang, L., 2021. The pineapple MADS-box gene family and the evolution of early monocot flower. Scientific reports, 11(1), pp.1–14.

Jiang, J., Ma, S., Ye, N., Jiang, M., Cao, J. and Zhang, J., 2017. WRKY transcription factors in plant responses to stresses. Journal of integrative plant biology, 59(2), pp.86–101.

Klosinska, M., Picard, C.L. and Gehring, M., 2016. Conserved imprinting associated with unique epigenetic signatures in the Arabidopsis genus. Nature Plants, 2(10), pp.1–8.

Lai, X., Blanc-Mathieu, R., Grandvuillemin, L., Huang, Y., Stigliani, A., Lucas, J., Thévenon, E., Loue-Manifel, J., Turchi, L., Daher, H. and Brun-Hernandez, E., 2021. The LEAFY floral regulator displays pioneer transcription factor properties. Molecular Plant, 14(5), pp.829–837.

Lee, D.S., Chen, L.J., Li, C.Y., Liu, Y., Tan, X.L., Lu, B.R., Li, J., Gan, S.X., Kang, S.G., Suh, H.S. and Zhu, Y., 2013. The Bsister MADS gene FST determines ovule patterning and development of the zygotic embryo and endosperm. PloS one, 8(3), p.e58748.

Li, G., Kuijer, H.N., Yang, X., Liu, H., Shen, C., Shi, J., Betts, N., Tucker, M.R., Liang, W., Waugh, R. and Burton, R.A., 2021. MADS1 maintains barley spike morphology at high ambient temperatures. Nature Plants, 7(8), pp.1093–1107.

Li, G., Liang, W., Zhang, X., Ren, H., Hu, J., Bennett, M.J. and Zhang, D., 2014. Rice actin-binding protein RMD is a key link in the auxin–actin regulatory loop that controls cell growth. Proceedings of the National Academy of Sciences, 111(28), pp.10377–10382.

Li, J., Yang, D.L., Huang, H., Zhang, G., He, L.I., Pang, J., Lozano-Durán, R., Lang, Z. and Zhu, J.K., 2020. Epigenetic memory marks determine epiallele stability at loci targeted by de novo DNA methylation. Nature Plants, 6(6), pp.661–674.

Li, X., Harris, C.J., Zhong, Z., Chen, W., Liu, R., Jia, B., Wang, Z., Li, S., Jacobsen, S.E. and Du, J., 2018. Mechanistic insights into plant SUVH family H3K9 methyltransferases and their binding to context-biased non-CG DNA methylation. Proceedings of the National Academy of Sciences, 115(37), pp.E8793–E8802.

Liu, C., Lu, F., Cui, X. and Cao, X., 2010. Histone methylation in higher plants. Annual review of plant biology, 61(1), pp.395–420.

Long, J., Walker, J., She, W., Aldridge, B., Gao, H., Deans, S., Vickers, M. and Feng, X., 2021. Nurse cell–derived small RNAs define paternal epigenetic inheritance in Arabidopsis. Science, 373(6550), p.eabh0556.

Lu, Jing, and Enrico Magnani. “Seed tissue and nutrient partitioning, a case for the nucellus.” Plant reproduction 31.3 (2018): 309–317.

Ma, X., Zhang, Q., Zhu, Q., Liu, W., Chen, Y., Qiu, R., Wang, B., Yang, Z., Li, H., Lin, Y. and Xie, Y., 2015. A robust CRISPR/Cas9 system for convenient, high-efficiency multiplex genome editing in monocot and dicot plants. Molecular plant, 8(8), pp.1274–1284.

Marchant, D.B. and Walbot, V., 2022. Anther development—The long road to making pollen. The Plant Cell, 34(12), pp.4677–4695.

Masiero, S., Imbriano, C., Ravasio, F., Favaro, R., Pelucchi, N., Gorla, M.S., Mantovani, R., Colombo, L. and Kater, M.M., 2002. Ternary complex formation between MADS-box transcription factors and the histone fold protein NF-YB. Journal of Biological Chemistry, 277(29), pp.26429–26435.

Mathieu, O., Probst, A.V. and Paszkowski, J., 2005. Distinct regulation of histone H3 methylation at lysines 27 and 9 by CpG methylation in Arabidopsis. The EMBO journal, 24(15), pp.2783–2791.

Mendes, M.A., Petrella, R., Cucinotta, M., Vignati, E., Gatti, S., Pinto, S.C., Bird, D.C., Gregis, V., Dickinson, H., Tucker, M.R. and Colombo, L., 2020. The RNA-dependent DNA methylation pathway is required to restrict SPOROCYTELESS/NOZZLE expression to specify a single female germ cell precursor in Arabidopsis. Development, 147(23), p.dev194274.

Mizzotti, C., Mendes, M.A., Caporali, E., Schnittger, A., Kater, M.M., Battaglia, R. and Colombo, L., 2012. The MADS box genes SEEDSTICK and ARABIDOPSIS Bsister play a maternal role in fertilization and seed development. The Plant Journal, 70(3), pp.409–420.

Nayar, S., Kapoor, M. and Kapoor, S., 2014. Post-translational regulation of rice MADS29 function: homodimerization or binary interactions with other seed-expressed MADS proteins modulate its translocation into the nucleus. Journal of experimental botany, 65(18), pp.5339–5350.

Nayar, S., Sharma, R., Tyagi, A.K. and Kapoor, S., 2013. Functional delineation of rice MADS29 reveals its role in embryo and endosperm development by affecting hormone homeostasis. Journal of Experimental Botany, 64(14), pp.4239–4253.

Nesi, N., Debeaujon, I., Jond, C., Stewart, A.J., Jenkins, G.I., Caboche, M. and Lepiniec, L., 2002. The TRANSPARENT TESTA16 locus encodes the ARABIDOPSIS BSISTER MADS domain protein and is required for proper development and pigmentation of the seed coat. The Plant Cell, 14(10), pp.2463–2479.

Ng, M. and Yanofsky, M.F., 2001. Function and evolution of the plant MADS-box gene family. Nature Reviews Genetics, 2(3), pp.186–195.

Nic-Can, G., Hernández-Castellano, S., Kú-González, A., Loyola-Vargas, V.M. and De-la-Peña, C., 2013. An efficient immunodetection method for histone modifications in plants. Plant Methods, 9(1), pp.1–9.

Nogueira, F.M., Nogueira, P.V.F., Vanzela, A.L.L. and Rocha, D.M., 2021. Ultrastructural analysis of Rhynchospora ovules: The first record of Cyperaceae megagametophyte on transmission electron microscope. Micron, 140, p.102962.

Oliver, C., Pradillo, M., Corredor, E. and Cuñado, N., 2013. The dynamics of histone H3 modifications is species-specific in plant meiosis. Planta, 238(1), pp.23–33.

Olmedo-Monfil, V., Durán-Figueroa, N., Arteaga-Vázquez, M., Demesa-Arévalo, E., Autran, D., Grimanelli, D., Slotkin, R.K., Martienssen, R.A. and Vielle-Calzada, J.P., 2010. Control of female gamete formation by a small RNA pathway in Arabidopsis. Nature, 464(7288), pp.628–632.

Parish, R.W. and Li, S.F., 2010. Death of a tapetum: a programme of developmental altruism. Plant Science, 178(2), pp.73–89.

Pinto, S.C., Mendes, M.A., Coimbra, S. and Tucker, M.R., 2019. Revisiting the female germline and its expanding toolbox. Trends in Plant Science, 24(5), pp.455–467.

Pinyopich, A., Ditta, G.S., Savidge, B., Liljegren, S.J., Baumann, E., Wisman, E. and Yanofsky, M.F., 2003. Assessing the redundancy of MADS-box genes during carpel and ovule development. Nature, 424(6944), pp.85–88.

Prasad, K., Zhang, X., Tobón, E. and Ambrose, B.A., 2010. The Arabidopsis B - sister MADS - box protein, GORDITA, represses fruit growth and contributes to integument development. The Plant Journal, 62(2), pp.203–214.

Radchuk, V., Borisjuk, L., Radchuk, R., Steinbiss, H.H., Rolletschek, H., Broeders, S. and Wobus, U., 2006. Jekyll encodes a novel protein involved in the sexual reproduction of barley. The Plant Cell, 18(7), pp.1652–1666.

Rodríguez-Leal, D., León-Martínez, G., Abad-Vivero, U. and Vielle-Calzada, J.P., 2015. Natural variation in epigenetic pathways affects the specification of female gamete precursors in Arabidopsis. The Plant Cell, 27(4), pp.1034–1045.

Rojek, J., Tucker, M.R., Rychłowski, M., Nowakowska, J. and Gutkowska, M., 2021. The Rab geranylgeranyl transferase beta subunit is essential for embryo and seed development in Arabidopsis thaliana. International journal of molecular sciences, 22(15), p.7907.

Rudall, P.J., 2021. Evolution and patterning of the ovule in seed plants. Biological Reviews, 96(3), pp.943–960.

Russell, S.D., 1979. Fine structure of megagametophyte development in Zea mays. Canadian Journal of Botany, 57(10), pp.1093–1110.

Schilling, S., Gramzow, L., Lobbes, D., Kirbis, A., Weilandt, L., Hoffmeier, A., Junker, A., Weigelt-Fischer, K., Klukas, C., Wu, F. and Meng, Z., 2015. Non - canonical structure, function and phylogeny of the B sister MADS- box gene O s MADS 30 of rice (O ryza sativa). The Plant Journal, 84(6), pp.1059–1072.

She, W., Grimanelli, D., Rutowicz, K., Whitehead, M.W., Puzio, M., Kotliński, M., Jerzmanowski, A. and Baroux, C., 2013. Chromatin reprogramming during the somatic-to-reproductive cell fate transition in plants. Development, 140(19), pp.4008–4019.

Shen, C.Y., Chen, Y.Y., Liu, K.W., Lu, H.C., Chang, S.B., Hsiao, Y.Y., Yang, F., Zhu, G., Zou, S.Q., Huang, L.Q. and Liu, Z.J., 2021. Orchid Bsister gene PeMADS28 displays conserved function in ovule integument development. Scientific reports, 11(1), pp.1–13.

Shoesmith, J.R., Solomon, C.U., Yang, X., Wilkinson, L.G., Sheldrick, S., van Eijden, E., Couwenberg, S., Pugh, L.M., Eskan, M., Stephens, J. and Barakate, A., 2021. APETALA2 functions as a temporal factor together with BLADE-ON-PETIOLE2 and MADS29 to control flower and grain development in barley. Development, 148(5), p.dev194894.

Singh, M., Goel, S., Meeley, R.B., Dantec, C., Parrinello, H., Michaud, C., Leblanc, O. and Grimanelli, D., 2011. Production of viable gametes without meiosis in maize deficient for an ARGONAUTE protein. The Plant Cell, 23(2), pp.443–458.

Su, Z., Wang, N., Hou, Z., Li, B., Li, D., Liu, Y., Cai, H., Qin, Y. and Chen, X., 2020. Regulation of female germline specification via small RNA mobility in Arabidopsis. Plant Cell, 32(9), pp.2842–2854.

Su, Z., Zhao, L., Zhao, Y., Li, S., Won, S., Cai, H., Wang, L., Li, Z., Chen, P., Qin, Y. and Chen, X., 2017. The THO complex non-cell-autonomously represses female germline specification through the TAS3-ARF3 module. Current Biology, 27(11), pp.1597–1609.

Waddington, S.R., Cartwright, P.M. and Wall, P.C., 1983. A quantitative scale of spike initial and pistil development in barley and wheat. Annals of Botany, 51(1), pp.119–130.

Wang, J., Guo, X., Xiao, Q., Zhu, J., Cheung, A.Y., Yuan, L., Vierling, E. and Xu, S., 2021. Auxin efflux controls orderly nucellar degeneration and expansion of the female gametophyte in Arabidopsis. New Phytologist, 230(6), pp.2261–2274.

Wang, Y., Ye, H., Bai, J. and Ren, F., 2020. The regulatory framework of developmentally programmed cell death in floral organs: A review. Plant Physiology and Biochemistry.

Wei, L., Gu, L., Song, X., Cui, X., Lu, Z., Zhou, M., Wang, L., Hu, F., Zhai, J., Meyers, B.C. and Cao, X., 2014. Dicer-like 3 produces transposable element-associated 24-nt siRNAs that control agricultural traits in rice. Proceedings of the National Academy of Sciences, 111(10), pp.3877–3882.

Wilkinson, L.G., 2019. Molecular and genetic cues influencing ovule development in barley (Hordeum vulgare) (Doctoral dissertation).

Wilkinson, L.G., Yang, X., Burton, R.A., Würschum, T. and Tucker, M.R., 2019. Natural variation in ovule morphology is influenced by multiple tissues and impacts downstream grain development in barley (Hordeum vulgare L.). Frontiers in plant science, 10, p.1374.

Xu, L., Yuan, K., Yuan, M., Meng, X., Chen, M., Wu, J., Li, J. and Qi, Y., 2020. Regulation of rice tillering by RNA-directed DNA methylation at miniature inverted-repeat transposable elements. Molecular plant, 13(6), pp.851–863.

Xu, W., Fiume, E., Coen, O., Pechoux, C., Lepiniec, L. and Magnani, E., 2016. Endosperm and nucellus develop antagonistically in Arabidopsis seeds. The Plant Cell, 28(6), pp.1343–1360.

Yang, F., Xu, F., Wang, X., Liao, Y., Chen, Q. and Meng, X., 2016. Characterization and functional analysis of a MADS-box transcription factor gene (GbMADS9) from Ginkgo biloba. Scientia Horticulturae, 212, pp.104–114.

Yang, W.C., Ye, D., Xu, J. and Sundaresan, V., 1999. The SPOROCYTELESS gene of Arabidopsis is required for initiation of sporogenesis and encodes a novel nuclear protein. Genes & Development, 13(16), pp.2108–2117.

Yang, X., Wilkinson, L.G., Aubert, M.K., Houston, K., Shirley, N., Tucker, M.R., 2022. Ovule cell wall composition is a maternal determinant of grain size in barley. bioRxiv

Yang, X., Wu, F., Lin, X., Du, X., Chong, K., Gramzow, L., Schilling, S., Becker, A., Theißen, G. and Meng, Z., 2012. Live and let die-the Bsister MADS-box gene OsMADS29 controls the degeneration of cells in maternal tissues during seed development of rice (Oryza sativa). PloS one, 7(12), p.e51435.

Yin, L.L. and Xue, H.W., 2012. The MADS29 transcription factor regulates the degradation of the nucellus and the nucellar projection during rice seed development. The Plant Cell, 24(3), pp.1049–1065.

Zhang, H., Lang, Z. and Zhu, J.K., 2018. Dynamics and function of DNA methylation in plants. Nature reviews Molecular cell biology, 19(8), pp.489–506.

Zhou, J.M. and Zhang, Y., 2020. Plant immunity: danger perception and signaling. Cell, 181(5), pp.978–989.

