## Extended Data for "MADS31 supports female germline development by repressing the post-fertilization program in cereal ovules"

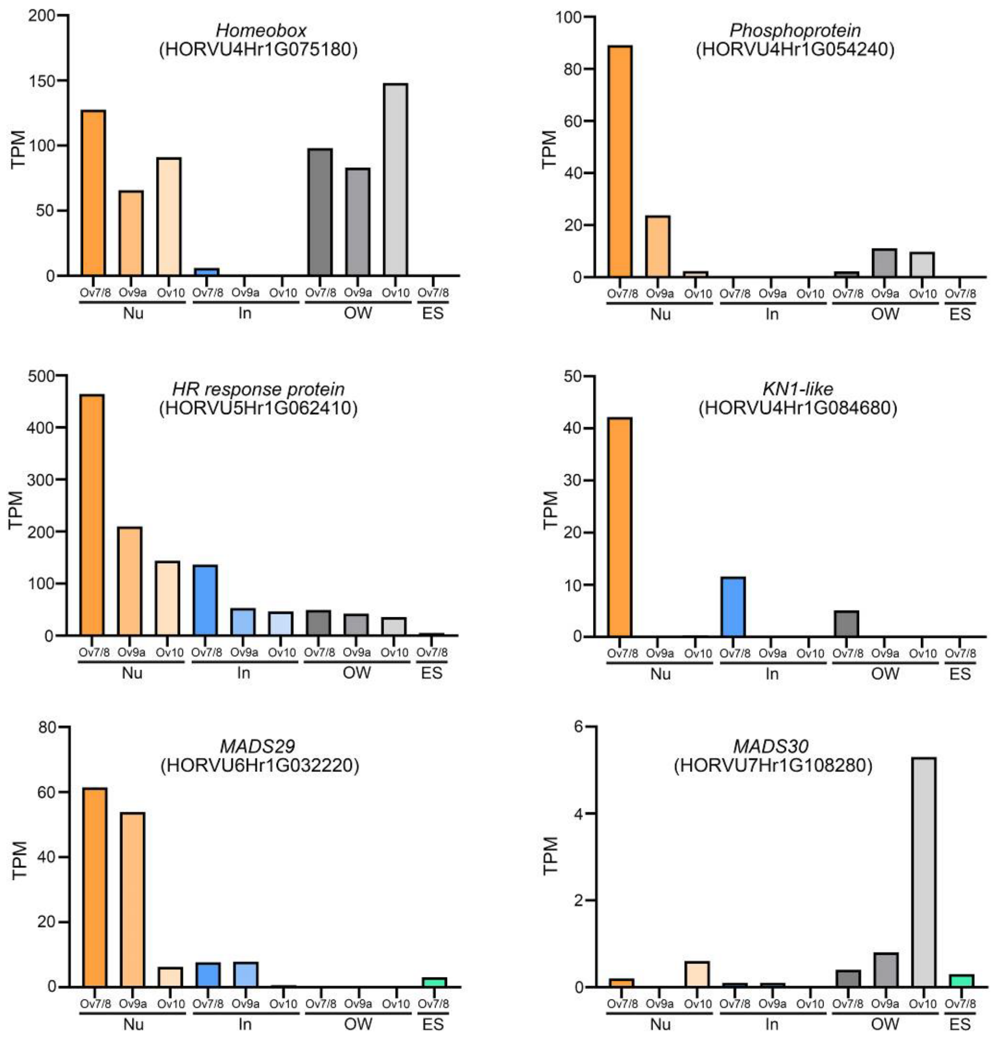


**Extended Data Fig. 1: Examples of genes expressed in the nucellus of the barley ovule.**
HR, heat response. KN1, KNOTTED1. MADS29 and MADS30 are the other two members of B sister class of MADS box genes.

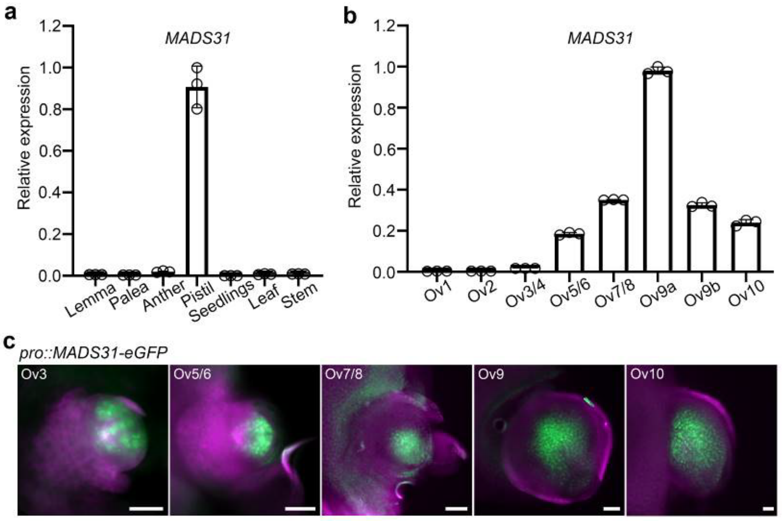


**Extended Data Fig. 2: Spatial and temporal expression pattern of *MADS31*.**
**a,** Relative expression level (qRT-PCR) of *MADS31* in vegetative tissues and floral organs. The data are shown as mean ± s.d.; *n* = 3 replicates. **b,** Relative expression level of *MADS31* during ovule development. Ov1, ovule primordium stage. Ov2, archesporial cell stage. Ov3/4, megaspore mother cell/meiosis stage. Ov5/6, functional megaspore stage. Ov7/8, female gametophyte mitosis stage. Ov9a, mature female gametophyte stage. Ov9b, embryo sac expansion stage. Ov10, anthesis stage. For Ov1–Ov3/4 stages, spikes were collected for RNA extraction. For Ov7/8–Ov10 stages, pistils were dissected from spikelets for RNA extraction. The data are shown as mean ± s.d.; *n* = 3 replicates. **c,** Accumulation of MADS31 protein during ovule development in *pro::MADS31-eGFP* transgenic lines. UV channel is used as background emission shown in magenta. Scale bars, 50 μm.

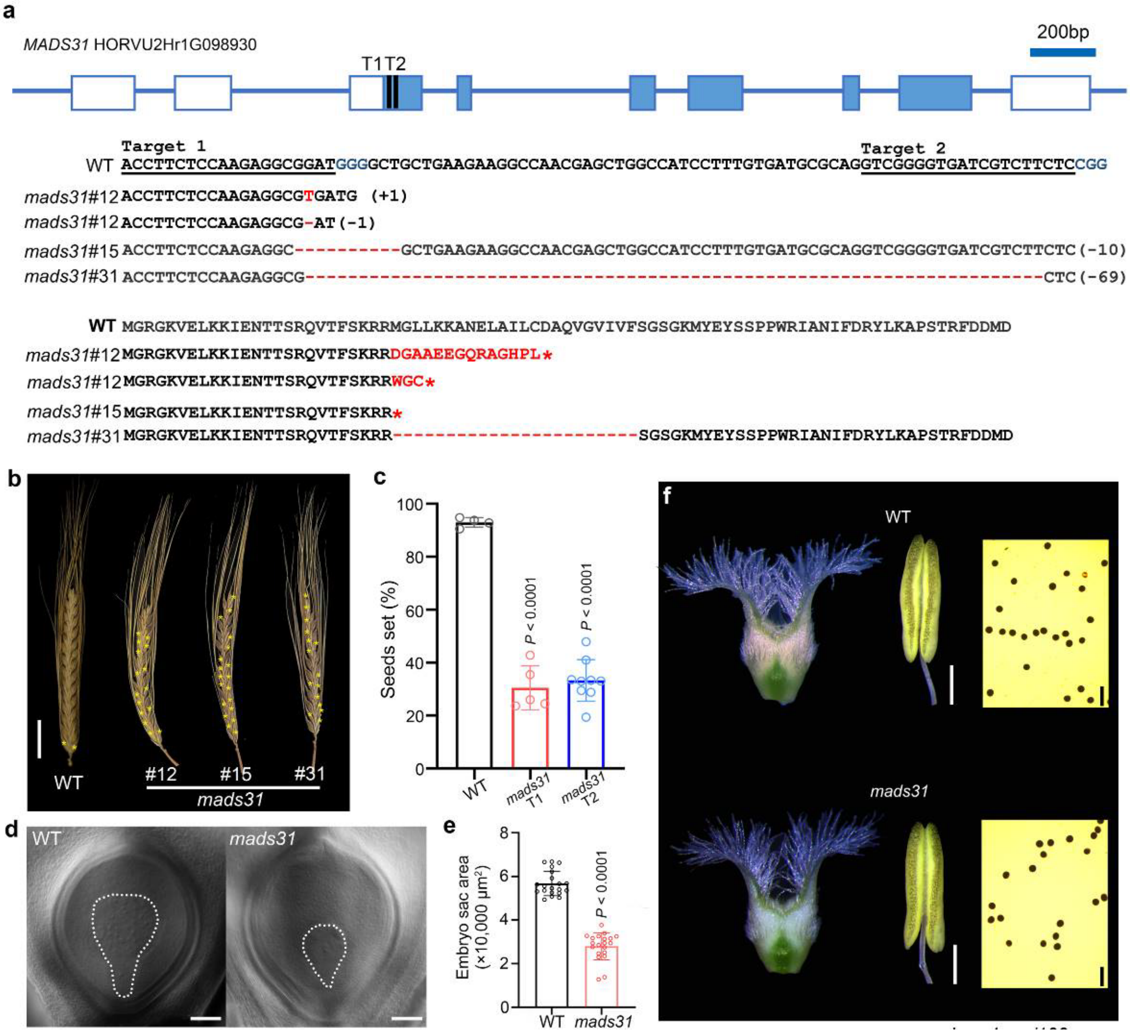


**Extended Data Fig. 3: Creation of barley *mads31* mutants using CRISPR/Cas9.**
**a,** Upper, the gene structure of *MADS31* and the positions of two sgRNA targets (T1 and T2) for gene editing. Blue boxes are exons and white boxes are UTRs. Middle, DNA sequences of independent T0 transgenic lines harboring mutations in *MADS31*. Lower, putative amino acid sequences in *mads31* mutants. Asterisks indicate a stop codon. WT, wild type. **b,** Mature spikes of WT and three alleles of *mads31*. Yellow asterisks indicate sterile spikelets. Scale bars, 2 cm. **c,** Seed set rate of WT and *mads31* (T1 and T2 generation transgenic lines). The data are shown as mean ± s.d.; 4, 5 and 9 individual plants are used for each genotype. **d,** Ovules from cleared pistils in WT and *mads31.* White dashed lines indicate embryo sacs. Scale bars, 100 μm. **e,** Measurement of embryo sac area from cleared pistils. The data are shown as mean ± s.d.; $n = 20$ replicates. **f,** Pistils and anthers of WT and *mads31*. Left, pistils at anthesis. Middle, anthers at anthesis. Scale bars, 1 mm. Right, pollen stained by $I_2$-KI solution. Scale bars, 100 μm.

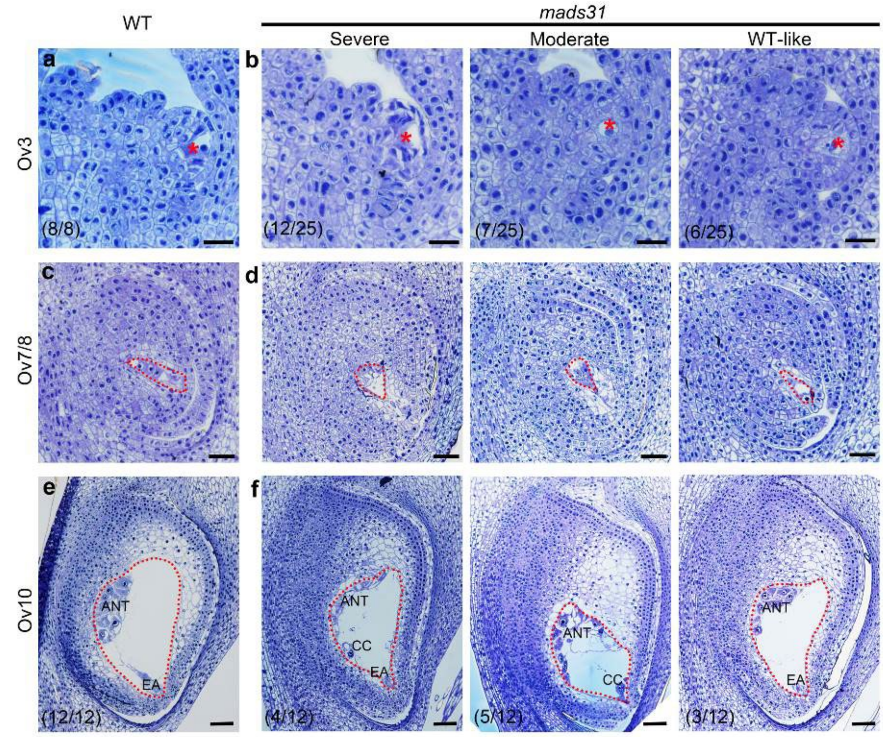


**Extended Data Fig. 4: Morphological changes in the inner nucellus and germline in *mads31.***
**a, b,** Longitudinal sections of wild-type (WT) and *mads31* ovules at Ov3 stage. Red asterisks indicate the megaspore mother cell. Scale bars, 25 μm. **c, d,** Longitudinal sections of WT and *mads31* ovules at Ov7/8 stage. Red dashed lines indicate the embryo sac. Scale bars, 50 μm. **e, f,** Longitudinal sections of WT and *mads31* ovules at Ov10 stage. Red dashed lines indicate the embryo sac. ANT, antipodal cells; CC, central cell; EA, egg apparatus. Numbers in parentheses indicate frequencies of defective ovules in all examined ovules. Scale bars, 50 μm.

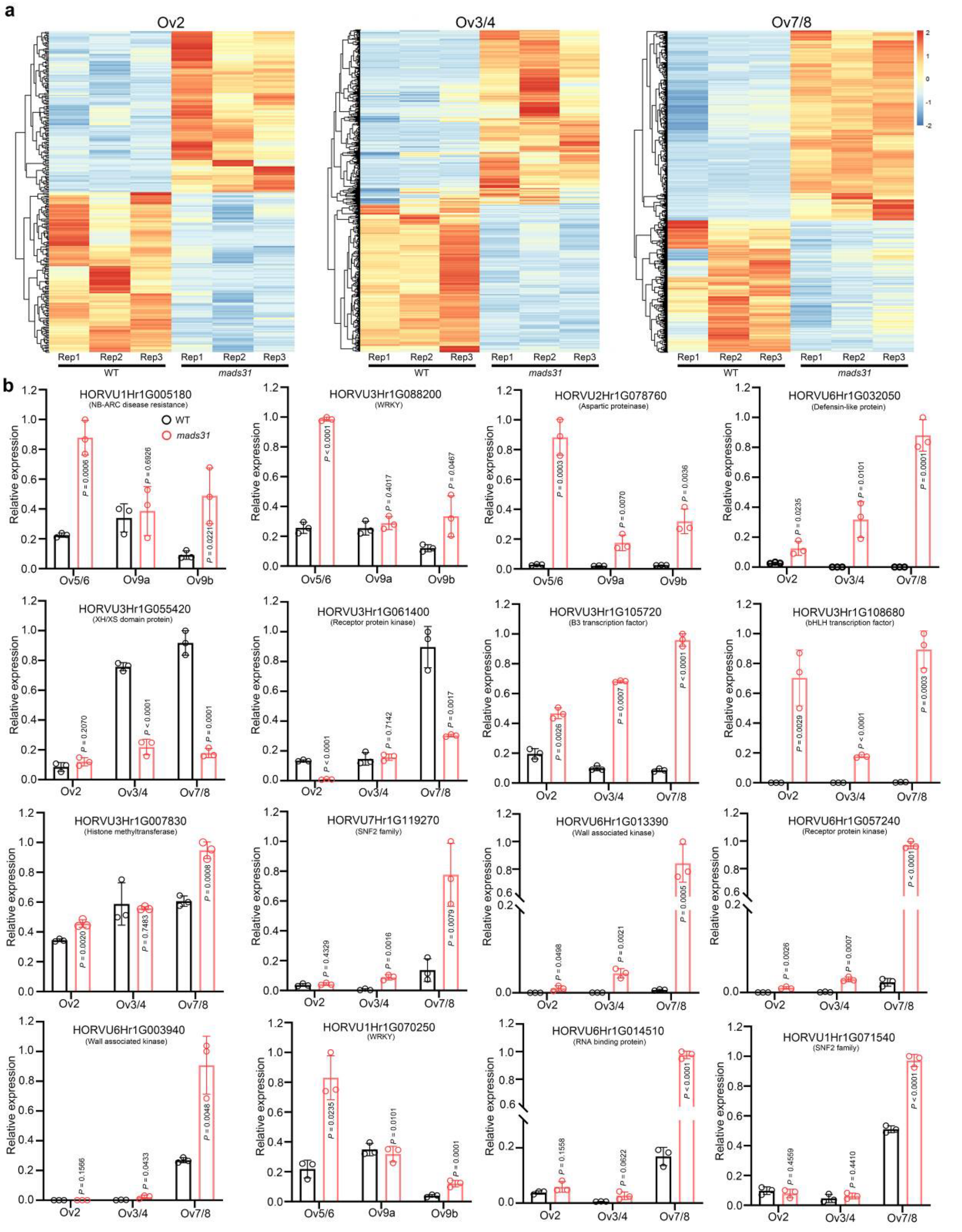


**Extended Data Fig. 5: Transcriptome profiling and verification of DEGs in *mads31*.**
**a,** Overview of differentially expressed genes (DEGs) between wild type (WT) and *mads31* at Ov2, Ov3, and Ov7/8 stages. **b,** Verifying expression change by qRT-PCR. The data are shown as mean ± s.d.; $n = 3$ replicates.

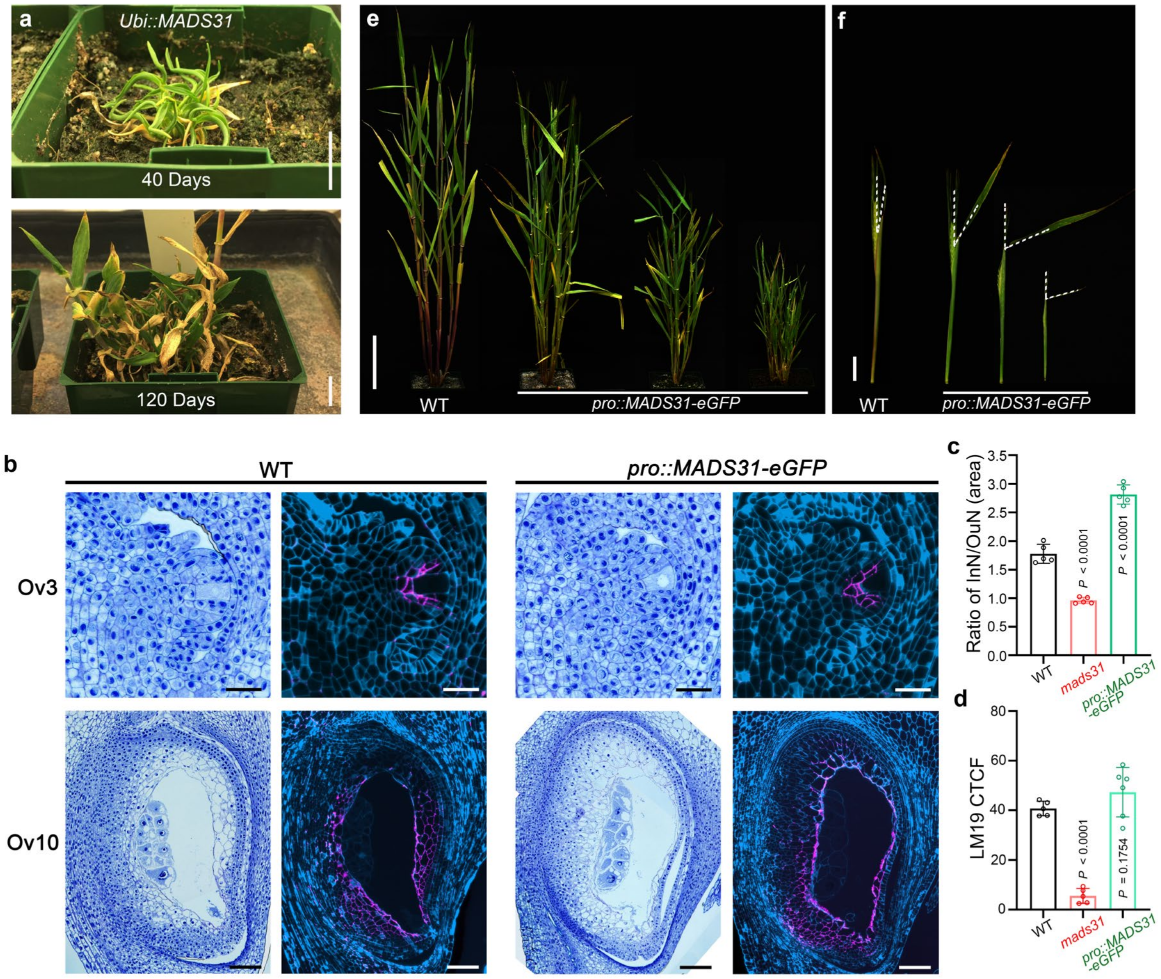


**Extended Data Fig. 6: Overexpression of *MADS31* represses plant growth.**
**a,** Extreme dwarfism in *Ubi::MADS31* transgenic plants. 40 days and 120 days indicate growth time after plants were transferred to soil. Scale bars, 1 cm. **b,** Nucellus patterning in wild-type (WT) and *pro::MADS31-eGFP* ovules. Left, toluidine blue stained longitudinal sections. Right, LM19 labeled demethylesterified pectin in the cell walls of nucellus. Scale bars, 25 μm (Ov3) and 100 μm (Ov10). **c,** Ratios of inner nucellus area versus outer nucellus area in WT, *mads31* and *pro::MADS31-eGFP* ovules. The data are shown as mean ± s.d.; $n = 5$ ovules. **d,** Corrected total cell fluorescence (CTCF) of LM19 immunosignals in the nucellus of WT, *mads31* and *pro::MADS31-eGFP* ovules. The data are shown as mean ± s.d.; $n = 5$ ovules. **e,** Various degrees of dwarfism in *pro::MADS31-eGFP* transgenic plants. Scale bar, 10 cm. **f,** Flag leaf inclination in wild-type (WT) and *pro::MADS31-eGFP* plants. Scale bar, 2 cm.

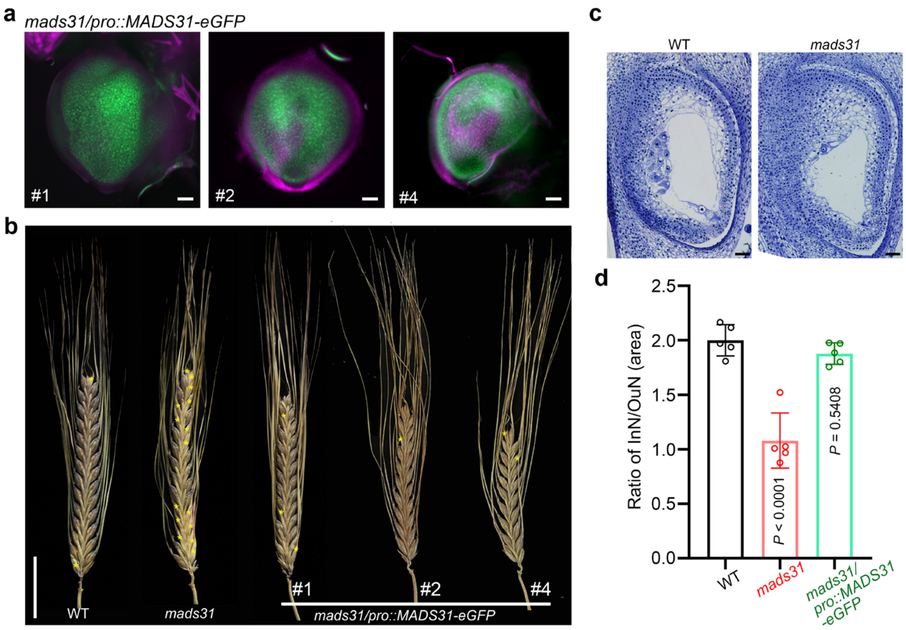


**Extended Data Fig. 7: Fusion of MADS31 and eGFP is able to rescue *mads31.***

**a,** Expression of MADS31-eGFP fusion protein in three transgenic *mads31* plants. Scale bars, 100 µm. **b,** Mature spikes of WT, *mads31* and *mads31/pro::MADS31-eGFP* plants. Yellow asterisks indicate sterile spikelets. Scale bar, 2 cm. **c,** Longitudinal sections of wild-type (WT), *mads31* and *mads31/pro::MADS31-eGFP* ovules. Scale bars, 50 µm. **d,** Ratios of inner nucellus area versus outer nucellus area in WT, *mads31* and mads31/*pro::MADS31-eGFP* ovules. The data are shown as mean ± s.d.; *n* = 5 ovules.

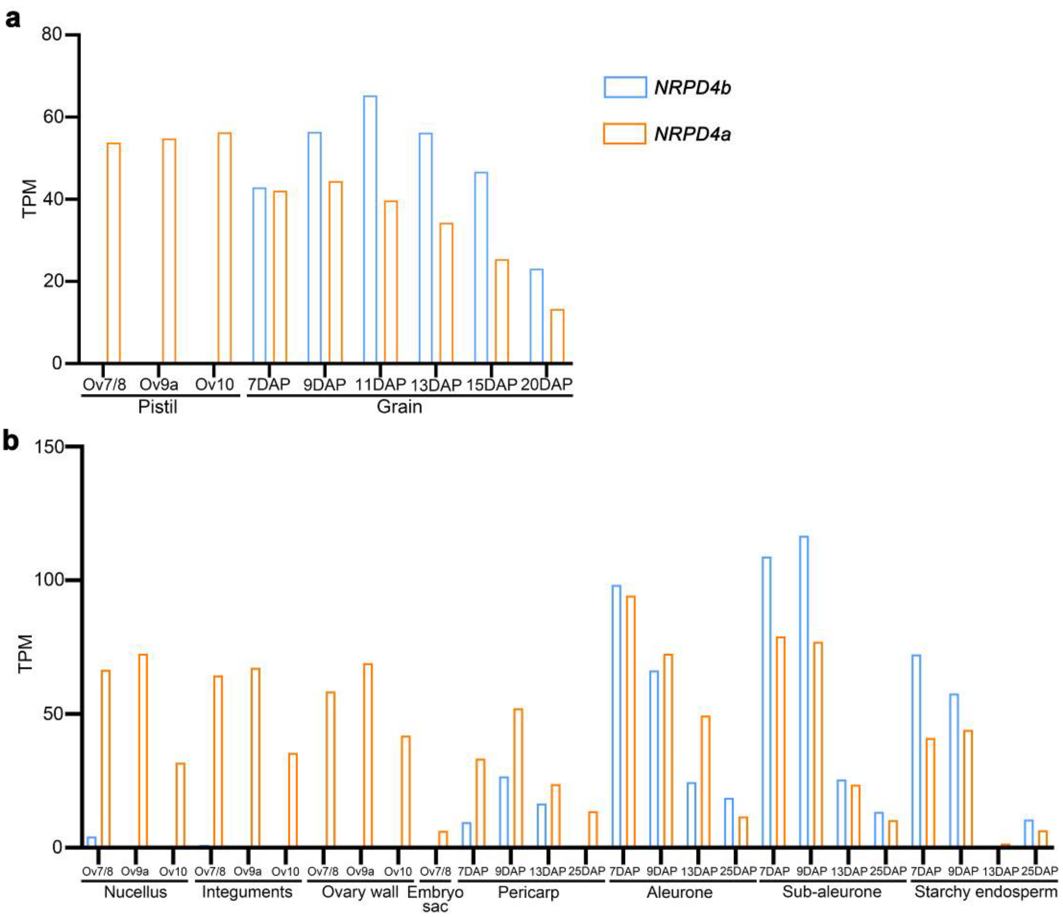


**Extended Data Fig. 8: Expression patterns of *NRPD4a* and *NRPD4b* in the pistil and grain.**
**a,** *NRPD4a and NRPD4b* expression in pistils and grains. **b,** *NRPD4a and NRPD4b* expression levels in various tissues of the pistil and grain. DAP, days after pollination. TPM, transcripts per million.

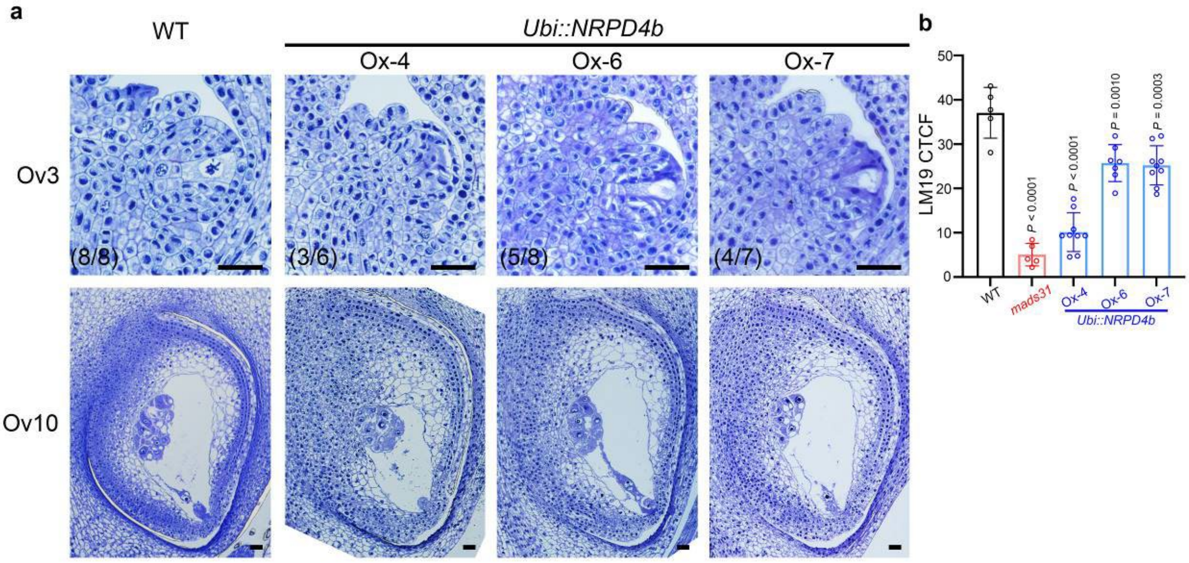


**Extended Data Fig. 9: Morphological changes of nucellus patterning in *Ubi::NRPD4b* lines.**
**a,** Longitudinal sections of wild-type (WT) and *Ubi::NRPD4b* ovules at Ov3 and Ov10 stages. Numbers in parentheses indicate frequencies of defective ovules in all examined ovules. Scale bars, 25 μm. **b,** Corrected total cell fluorescence (CTCF) of LM19 immunosignals in the nucellus of WT, *mads31* and *Ubi::NRPD4b* ovules. The data are shown as mean ± s.d.; *n* = 5–9 ovules.
